# Regulation of apical constriction via microtubule- and Rab11-dependent apical transport during tissue invagination

**DOI:** 10.1101/827378

**Authors:** Thao Phuong Le, SeYeon Chung

**Affiliations:** Department of Biological Sciences, Louisiana State University, Baton Rouge, LA 70803, USA

## Abstract

The formation of an epithelial tube is a fundamental process for organogenesis. During *Drosophila* embryonic salivary gland (SG) invagination, Folded gastrulation (Fog)- dependent Rho-associated kinase (Rok) promotes contractile apical myosin formation to drive apical constriction. Microtubules (MTs) are also crucial for this process and are required for forming and maintaining apicomedial myosin. However, the underlying mechanism that coordinates actomyosin and MT networks still remains elusive. Here, we show that MT-dependent intracellular trafficking regulates apical constriction during SG invagination. Key components involved in protein trafficking, such as Rab11 and Nuclear fallout (Nuf), are apically enriched near the SG invagination pit in a MT-dependent manner. Disruption of the MT networks or knockdown of *Rab11* impairs apicomedial myosin formation and apical constriction. We show that MTs and Rab11 are required for apical enrichment of the Fog ligand and the continuous distribution of the apical determinant protein Crumbs (Crb) and the key adherens junction protein E-Cadherin (E-Cad) along junctions. Targeted knockdown of *crb* or *E-Cad* in the SG disrupts apical myosin networks and results in apical constriction defects. Our data suggest a role of MT- and Rab11-dependent intracellular trafficking in regulating actomyosin networks and cell junctions, to coordinate cell behaviors during tubular organ formation.

## Introduction

Formation of three-dimensional tubes by invagination of flat epithelial sheets is a fundamental process in forming organs such as the lungs and kidneys (Andrew and Ewald, 2010). To enter the third dimension, cells must change their shapes and positions relative to each other. A major cellular process during epithelial tube formation is apical constriction, a universal cell shape change that is linked to tissue bending, folding and invagination (Sawyer *et al*., 2010; Martin and Goldstein, 2014). During apical constriction, the apical side of an epithelial cell constricts, causing a columnar or cuboidal cell to become wedge-shaped (Sawyer *et al*., 2010; Martin and Goldstein, 2014). Manipulation of apical constriction in a group of cells impacts both local and global tissue shape directly, suggesting a critical role for apical constriction in forming proper tissue architecture (Guglielmi *et al*., 2015; Chung *et al*., 2017; Izquierdo *et al*., 2018).

Apical constriction is driven by actin filament (F-actin) networks and the molecular motor non-muscle myosin II (hereafter referred to as myosin). Over the past decade, important functions of different actomyosin structures in epithelial tissue morphogenesis have been discovered. Particularly, studies in *Drosophila* revealed a distinct population of pulsatile apical medial actomyosin (hereafter referred to as apicomedial myosin) that generates a pulling force that exerts on adherens junctions to drive apical constriction (Martin *et al*., 2009; Rauzi *et al*., 2010; Booth *et al*., 2014; Chung *et al*., 2017). Further studies in early *Drosophila* embryos discovered that apicomedial myosin is created in response to signaling by the Folded gastrulation (Fog) ligand and its G protein-coupled receptors (GPCRs) (Manning *et al*., 2013; Kerridge *et al*., 2016) and is regulated by apical Rho-associated kinase (Rok) (Mason *et al*., 2013).

Apical junctions and apical determinants also have an important role in the formation of functional actomyosin complexes during developmental processes. During *Drosophila* dorsal closure, the apical polarity regulators Par-6, aPKC and Bazooka/Par3 (Baz/Par3) (altogether known as the Par complex) regulate pulsed actomyosin contractions in amnioserosa cells (David *et al*., 2010). In the *Drosophila* embryonic trachea, the apical protein Crumbs (Crb) is required for proper organization of the actomyosin complex (Letizia *et al*., 2011).

Emerging evidence suggests that microtubules (MTs) play a critical role in tissue invagination (Booth *et al*., 2014; Ko *et al*., 2019). MTs serve as tracks in intracellular transport (Le Droguen *et al*., 2015; Khanal *et al*., 2016; Aguilar-Aragon *et al*., 2020), raising the possibility that MTs regulate apical constriction through the endo- and exocytosis of membrane receptors and adhesion molecules. Several lines of evidence also suggest the importance of the endocytic pathway in apical constriction in both in vitro and in vivo systems. In *Drosophila* S2 cells, RhoGEF2 travels to the cell cortex by interaction with the MT plus-end protein EB1 to stimulate cell contraction (Rogers *et al*., 2004). During *Xenopus* gastrulation, disrupting endocytosis with dominant-negative dynamin or Rab5 perturbs apical constriction and invagination of cell sheets (Lee and Harland, 2010). During *Drosophila* gastrulation, the apical surface of cells is reshaped via Rab35 and RabGEF Sbf, which direct the plasma membrane to Rab11-positive recycling endosomes through a dynamic interaction with Rab5 endosomes to reshape actomyosin networks (Miao *et al*., 2019). Moreover, in the developing neural tube in *Xenopus*, asymmetric enrichment of Rab11 at the medial apical junctions is critical for apical constriction, suggesting that membrane trafficking has a key role in apical constriction (Ossipova *et al*., 2014). However, exactly how vesicle trafficking, MTs, and actomyosin networks are linked during tissue invagination remains to be discovered.

To determine how these three cellular attributes contribute to tissue invagination, we use the *Drosophila* embryonic SG as a model. We and others showed that apical constriction is regulated in a highly coordinated and spatiotemporally controlled manner during SG tube formation (Myat and Andrew, 2000; Booth *et al*., 2014; Chung *et al*., 2017; Sanchez-Corrales *et al*., 2018). The *Drosophila* embryo forms two SG tubes via invagination of two epithelial placodes on the ventral surface (Myat and Andrew, 2002; Chung *et al*., 2014). Before invagination, a small group of cells in the dorsal/posterior region of each SG placode begin to constrict. As those cells internalize to form the invagination pit, more cells anterior to the pit undergo clustered apical constriction in a Fog signaling-dependent manner (Chung *et al*., 2017) (Figure 1A). In the absence of *fog*, SG cells fail to accumulate Rok and myosin in the apicomedial region of the cells (Chung *et al*., 2017). MTs aid in forming and maintaining the apicomedial myosin network during SG invagination (Booth *et al*., 2014). The MT cytoskeleton near the invagination pit forms a network of longitudinal MT bundles, with the minus ends of MTs facing the apical domain of the cells and interacting with the apicomedial myosin (Booth *et al*., 2014). Disruption of MTs causes loss of apicomedial myosin and disrupted apical constriction during SG invagination (Booth *et al*., 2014). However, it is still unknown how intracellular trafficking affects these two processes.

**Figure 1.**
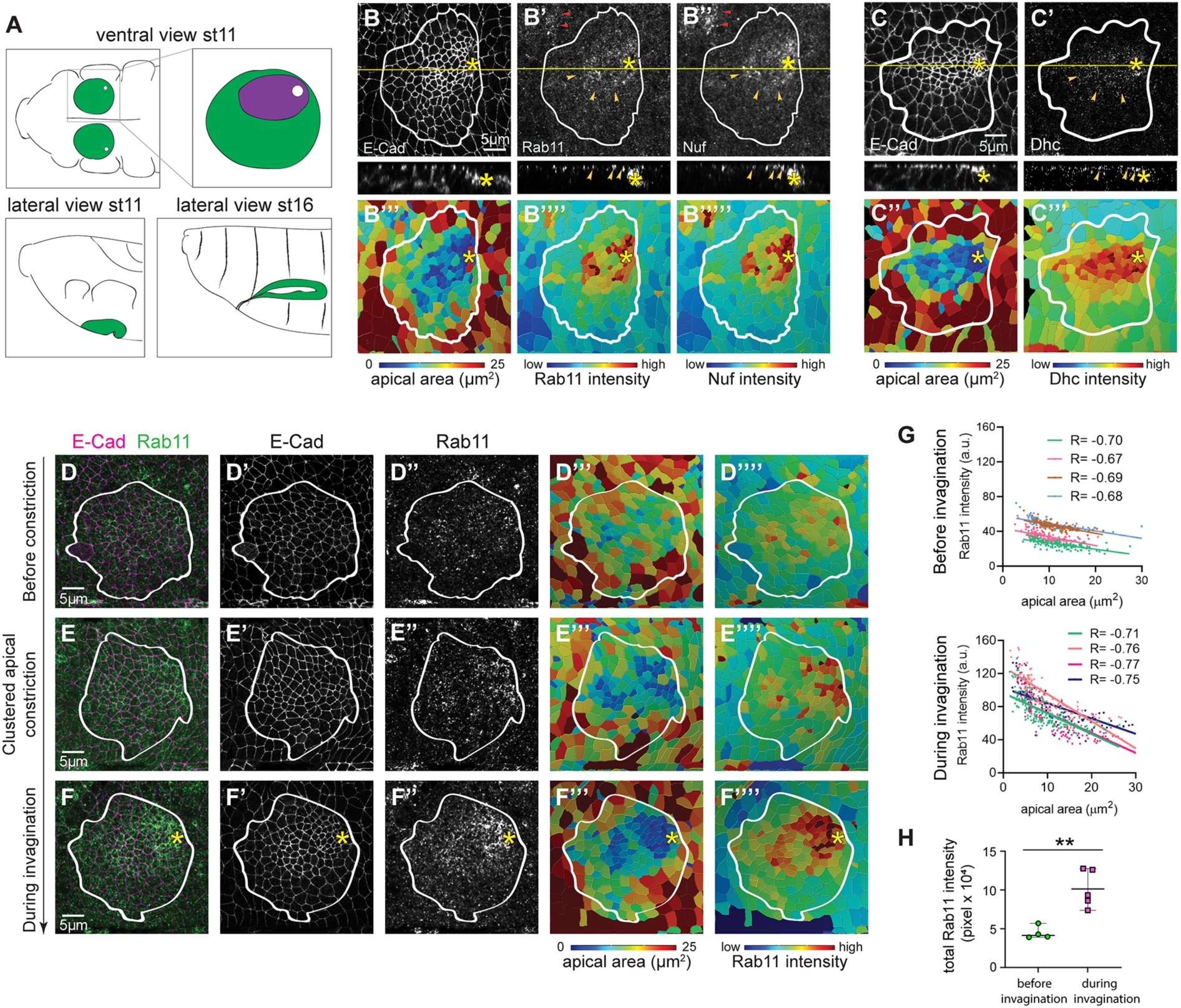
Intracellular trafficking components are apically enriched in invaginating SGs. (A) A schematic drawing of the anterior region of the *Drosophila* embryo for stage 11 (ventral and lateral views) and stage 16 (lateral view) with SGs shown in green. Top right, magnified view of a stage 11 SG. The region where SG cells undergo clustered apical constriction during invagination is shown in purple. (B-B’’) En face (top) and lateral (bottom) views of a wild type stage 11 SG immunostained for E-Cad, Rab11 and Nuf. Rab11 and Nuf show apical enrichment near the invagination pit (yellow arrowheads). Red arrowheads, Rab11 and Nuf signals near the segmental groove. (B’’’-B’’’’’) Heat maps of apical area (B’’’) and intensity of Rab11 (B’’’’) and Nuf (B’’’’’). Cells with smaller apical areas near the invagination pit (dark blue cells in B’’’) show the high intensity of Rab11 and Nuf (red signals in B’’’’ and B’’’’’). (C-C’’’) En face (top) and lateral (bottom) views of a wild type stage 11 SG immunostained for E-Cad (C) and Dhc64C (C’). Corresponding heat maps for apical areas (C’’) and intensity of Dhc64C signals (C’’’) are shown. (D-F’’’) Wild type stage 11 SGs immunostained for E-Cad and Rab11 at different timepoints of invagination. (G) The total intensity of Rab11 in the whole SG placode before and during invagination. Before invagination, n= 4 SGs; during invagination, n= 5 SGs. (H) Negative correlation between Rab11 intensities and apical areas of SG cells before (top) and during invagination (bottom). R, Pearson correlation coefficient. P< 0.0001 for all samples. Before invagination, n= 4 SGs, 506 cells; during invagination, n= 4 SGs, 561 cells. Asterisks: invagination pit. White lines: SG boundary. SG boundaries are marked based on CrebA (SG nuclear marker) signals in the basal region of SG cells (not shown).

In this study, we demonstrate that key proteins involved in intracellular trafficking - including Rab11, its binding partner Nuclear fallout (Nuf), and dynein heavy chain - are enriched in the apical domain of SG cells during invagination. Moreover, disruption of MTs results in mislocalization of Rab11 to the basolateral region of SG cells. Reducing *Rab11* in the SG leads to a decrease and dispersal of Rok and myosin in the apical domain and causes uncoordinated apical constriction. Our data suggest that apical localization of the Fog ligand, the apical determinant protein Crb, and the key adherens junction protein E-Cadherin (E-Cad) is compromised when the MT networks are disrupted or *Rab11* is knocked down. We further show that reducing *crb* or *E-Cad* in the SG leads to defects that are reminiscent of *Rab11* knockdown. Altogether, our work mechanistically links MT- and Rab11-dependent intracellular trafficking to the regulation of actomyosin networks during tubular organ formation.

## Results

### Intracellular trafficking components are apically enriched near the invagination pit

To test a role for the intracellular trafficking machinery in spatially biased signaling activation and protein accumulation during SG invagination, we analyzed the subcellular localization of proteins involved in vesicle trafficking in the SG. Several endosome markers and their interacting partners were tested, including Rab5 (an early endosome marker) (Gorvel *et al*., 1991), Rab7 (a late endosome marker) (Wichmann *et al*., 1992; Meresse *et al*., 1995), Rab11 (a recycling endosome marker) (Ullrich *et al*., 1996), Nuclear fallout (Nuf, a putative binding partner for Rab11) (Riggs *et al*., 2003), and Sec15 (an exocyst complex component and effector for Rab11) (Zhang *et al*., 2004; Langevin *et al*., 2005). Using labeling with E-Cad, an adherens junction marker, and CrebA, a SG nuclear marker, we segmented apical cell outlines and calculated the intensity mean of fluorescence signals of each endosomal marker. We observed apical enrichment of several of them in the SG, including Rab11 and Nuf (Figure 1, B-B’’’’’). Similar enrichment of Rab11 signals was also observed in the endogenously tagged Rab11 protein (Rab11-EYFP; (Dunst *et al*., 2015) (Supplemental Figure S1, A-A’’’’’). Importantly, the enrichment of Rab11 was more pronounced in the SG over time (Figure 1, D-F’’’’). Before invagination, only low levels of Rab11 were detected in the apical domain of all SG cells throughout the entire SG placode (Figure 1, D-D’’’’). Stronger signals of Rab11 were soon detected in the SG as cells begin to undergo apical constriction in the dorsal/posterior region of the placode (Figure 1, E-E’’’’), which intensified as invagination proceeded (Figure 1, F-F’’’’). Quantification of the total intensity of Rab11 in the whole SG placode showed significantly higher levels of Rab11 in invaginating SGs compared to before invagination (Figure 1H), suggesting upregulated Rab11 levels in SG cells during invagination. Furthermore, compared to before invagination (Figure 1G, top), intensity mean of Rab11 in the apical domain of each cell showed a stronger negative correlation with the apical area of cells in the entire SG during invagination (Figure 1G, bottom), suggesting a close link between intracellular trafficking activities and apical constriction.

EYFP-tagged Rab5, an early endosome marker, was also enriched in the apical domain in the invaginating SG, suggesting active endocytosis in the apical domain during SG invagination (Supplemental Figure S1, B-B’’’). Rab7, a late endosome marker, however, did not show apical enrichment but localized as large punctate structures in the cytoplasm of the entire SG placode (Supplemental Figure 1, D-D’’’). Dynein heavy chain 64C, a subunit of the dynein motor complex that transports cargos along MTs toward their minus ends, also showed apical enrichment (Figure 1, C-C’’’). This is consistent with the previous report that the minus end of MTs faces the apical domain of SG cells near the invagination pit (Booth *et al*., 2014). Sec15, an exocyst complex component, also showed similar apical enrichment (Supplemental Figure S1, E-E’’’). Similar to Rab11 (Figure 1G), intensity mean of Rab5-EYFP and Sec15 in the apical domain of each cell showed a negative correlation with the apical area of cells in the entire SG during invagination (Supplemental Figure S1, C and F). Compared to a more uniform distribution of other endosomal markers in the entire apical domain of SG cells, Sec15 signals appeared to be enriched at adherens junctions (Supplemental Figure S1E’). Overall, our results suggest active intracellular trafficking, possibly both endo- and exocytosis, in the apical domain of SG cells near the invagination pit during invagination.

### Apical enrichment of Rab11 and Nuf is MT-dependent

Strong apical enrichment of Rab11 and other trafficking components in SG cells near the invagination pit led us to test whether these vesicle markers are associated with vertically aligned MTs that could facilitate their polarized distribution. As in many epithelial cells, MT minus- and plus-ends face the apical and the basal domain of the SG cells, respectively (Myat and Andrew, 2002; Booth *et al*., 2014). Indeed, co-immunostaining of tyrosinated α-tubulin, a marker of dynamic or newly polymerized MTs (Westermann and Weber, 2003), and Rab11 showed a partial overlap of the two proteins at the apical region of the SG cells (Figure 2, A-B’’).

**Figure 2.**
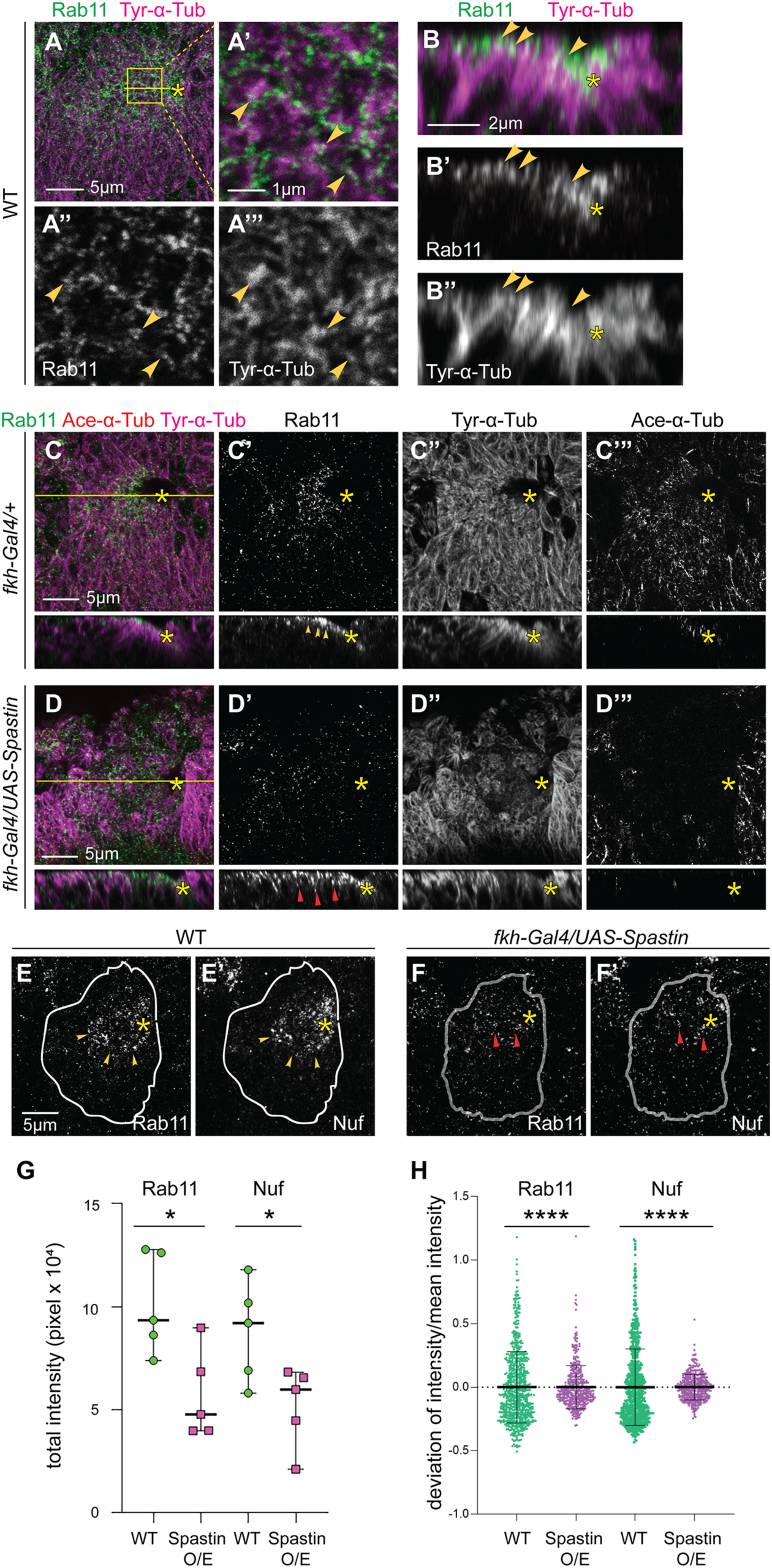
Disruption of MTs results in mislocalization of Rab11 and Nuf to the basolateral domain in SG cells. (A-A’’’) A wild type SG immunostained for Rab11 (green) and tyrosinated α-tubulin (Tyr-α-Tub; purple). Higher magnification of the yellow boxed area is shown in A’-A’’’. Yellow arrowheads, co-localized Rab11 and tyrosinated α-tubulin. (B-B’’) Z sections along the yellow line in A. (C-C’’’) En face (top) and lateral (bottom) views of a control (*fkh-Gal4/+*) SG show abundant levels of tyrosinated α-tubulin (C’’; Tyr-α-Tub) and acetylated α-tubulin (C’’’; Ace-α-Tub). (D-D’’’) Overexpression of spastin in the SG using *fkh-Gal4* leads to a reduction of tyrosinated α-tubulin (D’’) and loss of acetylated α-tubulin (D’’’). Whereas Rab11 is enriched in the apical domain of WT SG cells (yellow arrowheads in C’), Rab11 mislocalizes basolaterally when spastin is overexpressed (red arrowheads in D’). (E-F’’) Confocal images of WT (E and E’) and spastin-overexpressing SGs (F, F’) stained for Rab11 (E, F) and Nuf (E’, F’). The maximum projection of three confocal sections for the apical domain is shown. Compared to strong Rab11 and Nuf signals in the apical domain in WT SGs (yellow arrowheads in E and E’), Rab11 and Nuf are reduced in spastin-overexpressing SGs (red arrowheads in F and F’). (G) The total intensity of Rab11 and Nuf in the whole SG placode in control and spastin-overexpressing SG. n= 5 SGs for each genotype. (H) The degree of variability of Rab11 (left) and Nuf (right) intensities in the SG (n= 5 SGs for both genotypes; WT, 690 cells; *fkh-Gal4/UAS-Spastin*, 496 cells). ****, p<0.0001, Mann-Whitney U test. Asterisks: invagination pit. White lines: SG boundary.

To test whether Rab11 apical enrichment in SG cells is dependent on MT networks, we disrupted MTs in the SG by overexpressing spastin, a MT-severing protein (Sherwood *et al*., 2004), using the SG-specific *fkh-Gal4* driver. In control SGs, tyrosinated α-tubulin and acetylated α-tubulin, a marker of stable and longer-lived MTs (Westermann and Weber, 2003), were observed abundantly in the apical domain of cells in the whole placode (Figure 2, C’’ and C’’’; Supplemental Figure S2A’’). In spastin-overexpressing SGs, both tyrosinated and acetylated α-tubulin signals were strongly reduced, revealing a loss of MT filaments (Booth *et al*., 2014) (Figure 2, D’’ and D’’’ and Supplemental Figure S2B’’). Compared to control, spastin-overexpressing SGs showed cells with extremely small or large apical areas (Supplemental Figure S2, C-E). Cells with small apical areas were distributed randomly throughout the SG placode, rather than clustered as in control SGs (Supplemental Figure S2, C and D). The apical enrichment of Rab11 (Figure 2C’) was also disrupted and Rab11 was mislocalized basolaterally (Figure 2D’). SGs with disrupted MTs showed much lower Rab11 and Nuf intensity in the apical domain compared to control SGs (Figure 2, E-G). To compare the relative variability of apical enrichment of Rab11/Nuf in WT and spastin-overexpressing SGs, we calculated the degree of variability, the ratio of the deviation of Rab11/Nuf intensity to the mean Rab11/Nuf intensity. Compared to WT, spastin-overexpressing SGs showed a lower degree of relative variability of intensity (Figure 2G). These data suggest that apical Rab11/Nuf signals are less varied from cell to cell regardless of apical areas of individual cells in spastin-overexpressing SGs. Overall, these data suggest that MT networks are required for apical enrichment of Rab11 and Nuf during SG invagination.

### Rab11 and dynein functions are required for apical constriction in the SG

We next asked if modulation of Rab11 levels could compromise apical constriction. To test this possibility, we disrupted the function of Rab11 in the SG by either overexpressing a dominant-negative form of Rab11 (Rab11S25N-YFP; (Zhang *et al*., 2007); hereafter referred to as Rab11-DN) or knocking down *Rab11* using RNAi lines. Reduced Rab11 levels upon *Rab11* knockdown were confirmed using a Rab11 antibody in both stage 11 and stage 13 SGs (Supplemental Figure S3, A-D’). SGs that were invaginated within the range of 5.1-9.9 μm depth were used for quantification for proper comparison between different genotypes. Compared to control (Figure 3A), SGs overexpressing Rab11-DN or knocking down *Rab11* showed more cells with larger apical areas (Figure 3, B, C and F), suggesting a role for Rab11 in apical constriction during SG invagination.

**Figure 3.**
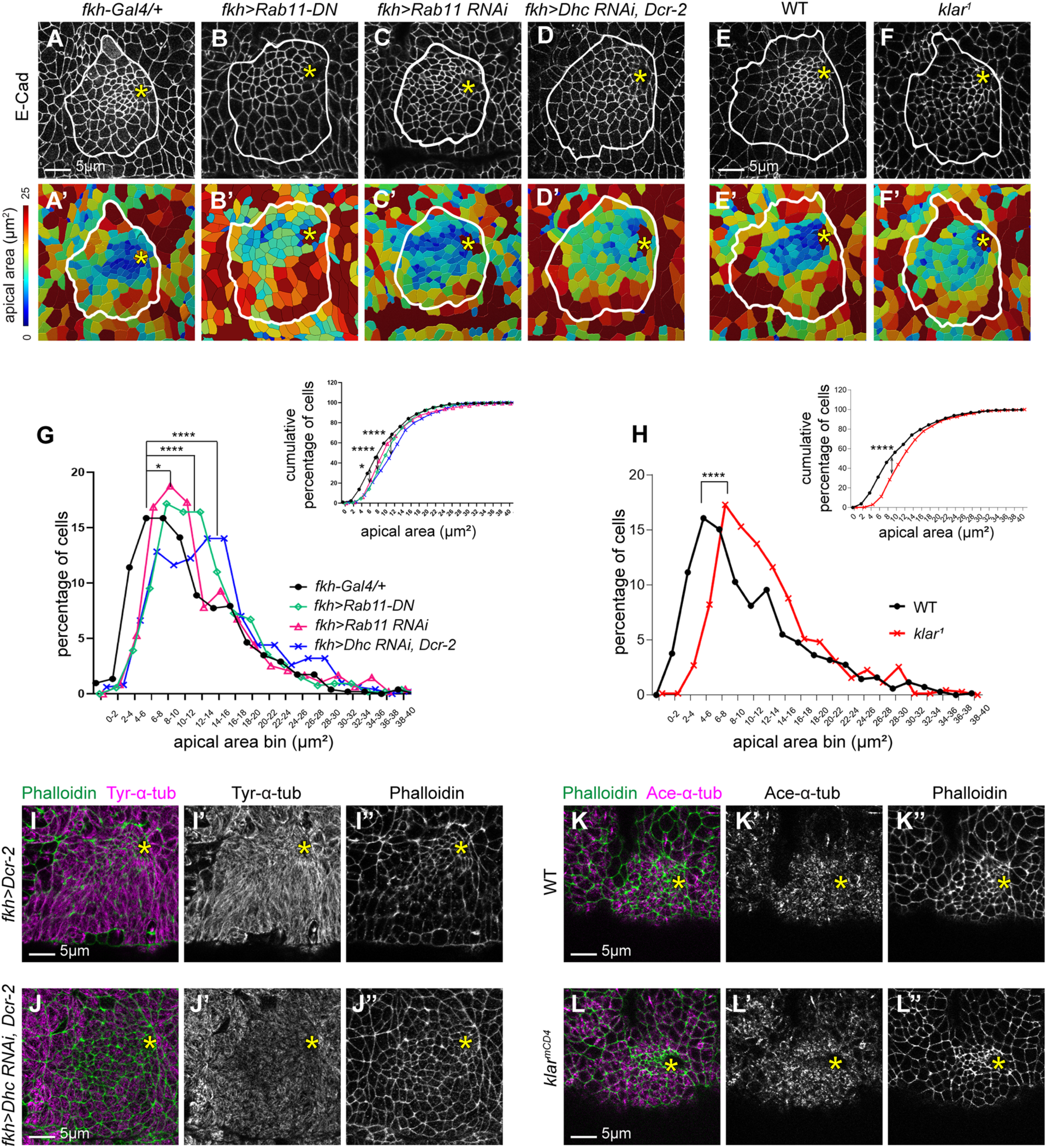
Rab11 and dynein functions are required for coordinated apical constriction in the SG. (A-F) Confocal images of control (A), Rab11-DN-overexpressing (B), *Rab11* RNAi (C), *Dhc64C* RNAi (D), wild type (E) and *klar* mutant (F) SGs immunostained for E-Cad. (A’-F’) Heat maps corresponding to images shown in A-F. (G, H) Percentage (G) and cumulative percentage (H) of SG cells with different apical areas. Mann-Whitney U test (for the percentage of cells) and Kolmogorov-Smirnov test (for the cumulative percentage of cells). N= 5 SGs (control (*fkh-Gal4/+*), 517 cells; *fkh>Rab11-DN*, 536 cells; *fkh>Rab11* RNAi, 474 cells; *fkh>Dhc64C* RNAi, 499 cells) and 6 SGs (WT, 690 cells; *klar^1^*, 705 cells). *, p<0.05; ****, p<0.0001. (I-J’’) Control (I-I’’) and *Dhc64C* RNAi (J-J’’) SGs stained for tyrosinated-*α*-Tub (I’, J’) and phalloidin (I’’, J’’). Compared to control (I’), tyrosinated-*α*-Tub (Tyr-*α*-Tub) is decreased in *Dhc64C* RNAi SGs (J’). (I-J’’) WT (K-K’’) and *klar* mutant (L-L’’) SGs stained for acetylated-*α*-Tub (Ace-*α*-Tub; K’, K’) and phalloidin (L’’, L’’). No significant changes in acetylated-*α*-Tub levels are detected in *klar* mutant SGs. Asterisks: invagination pit. White lines: SG boundary.

As dynein heavy chain was also enriched apically in the SG during invagination (Figure 1, C’ and C’’’), we tested if dynein function is also required for apical constriction during SG invagination. Indeed, knockdown of *Dynein heavy chain 64C* (*Dhc64C*) in the SG using RNAi resulted in more cells with larger apical areas during invagination, suggesting defective apical constriction upon *Dhc64C* knockdown (Figure 3, D, D’ and G). As dynein binds to and clusters the minus ends of microtubules (Heald *et al*., 1996; Khodjakov *et al*., 2003; Goshima *et al*., 2005; Burbank *et al*., 2006; Elting *et al*., 2014; Tan *et al*., 2018), we asked if *Dhc64C* knockdown affects the MT networks in the SG. Knocking down *Dhc64C* in the SG resulted in a slight reduction of tyrosinated α-tubulin (Figure 3J’). These data suggest that apical constriction defects observed in *Dhc64C* knockdown are, at least in part, due to affected MT networks.

Klarsicht (Klar), the *Drosophila* Klarsicht-Anc-Syne Homology (KASH) domain protein, mediates apical transport in the SG (Myat and Andrew, 2002) via the MT motor cytoplasmic dynein (Gross *et al*., 2000). We therefore tested if *klar* also influences apical constriction during SG invagination. *klar* null mutant embryos showed SGs with mild apical constriction defects compared to wild type (Figure 3, E-F’ and H). Importantly, unlike *Dhc64C* knockdown, no significant difference in the MT networks was observed in SGs in *klar* mutants (Figure 3L’), suggesting that apical constriction defects in *klar* mutants are due to disrupted apical transport. Overall, our data suggest essential roles of MT-, Rab11- and dynein-dependent intracellular transport in regulating apical constriction during SG invagination.

### Reduced Rab11 function leads to the reduction of apicomedial myosin formation and failure in accumulation of apicomedial Rok in SG cells

The apicomedial myosin structure generates the pulling force to drive apical constriction in SG cells (Booth *et al*., 2014; Chung *et al*., 2017). Disruption of MTs by spastin overexpression inhibits formation of apicomedial myosin during SG invagination (Booth *et al*., 2014). To test whether apicomedial myosin is affected in SG cells when *Rab11* is knocked down, we measured the overall intensity of myosin in the apicomedial region of SG cells using sqh-GFP, a functional GFP-tagged version of the myosin regulatory light chain (Royou *et al*., 2004). In control SGs, sqh-GFP showed strong myosin signals with clear web-like structures in the apicomedial region of cells near the invagination pit (Figure 4, A-A’’’’). Knockdown of *Rab11*, however, caused a significant reduction of apicomedial myosin intensity in SG cells in the same area (Figure 4, B-B’’’’ and C). Moreover, unlike clear web-like structures of apicomedial myosin in control SGs (Figure 4, A’’-A’’’’), myosin was more dispersed and fragmented in *Rab11* RNAi SGs (Figure 4, B’’-B’’’’), resulting in a decrease in areas of myosin accumulation in the apicomedial region of SG cells (Figure 4E). These data suggest roles for Rab11 in both upregulating apical myosin and forming and/or maintaining apicomedial myosin web.

**Figure 4.**
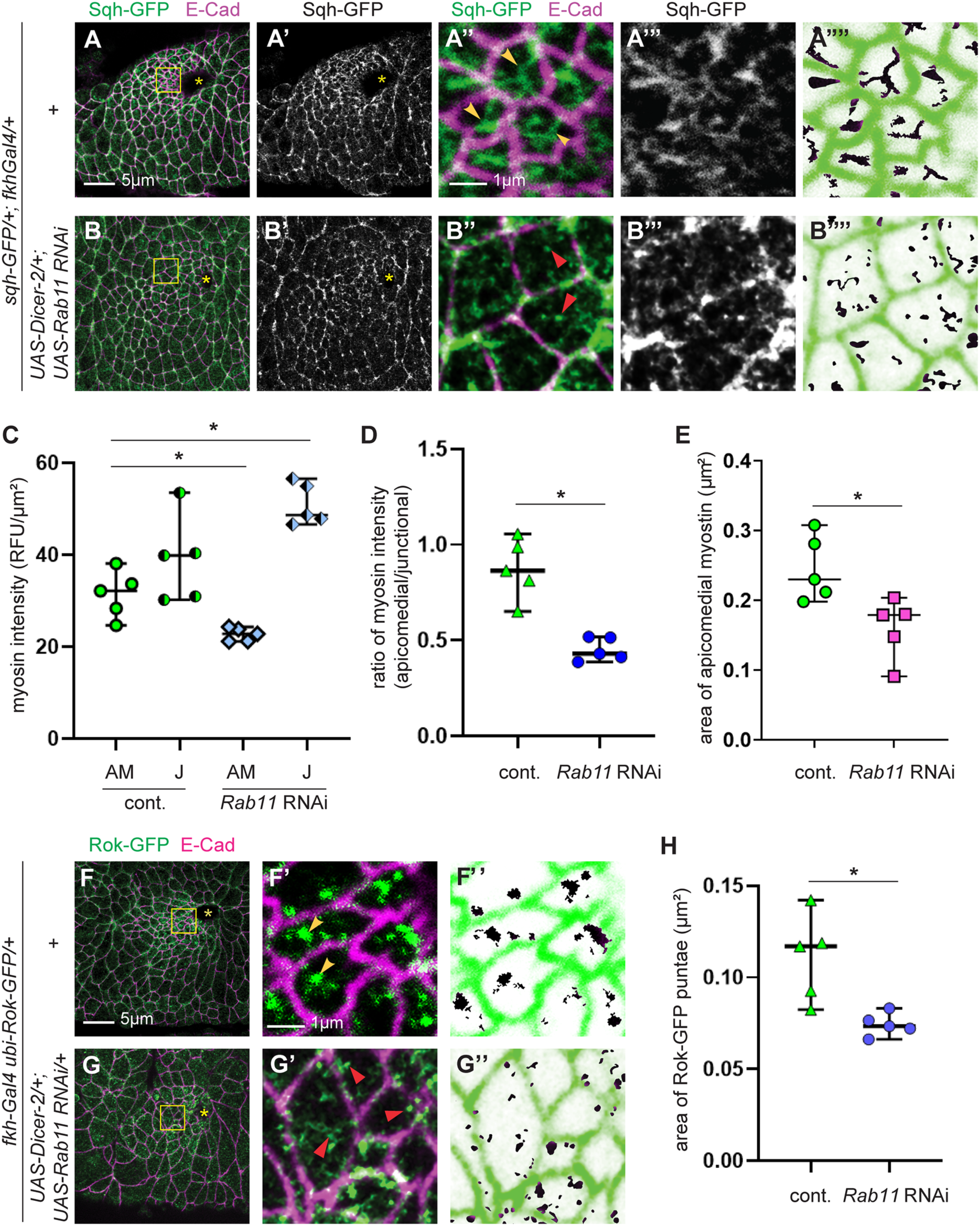
Compromised Rab11 functions lead to a reduction of apicomedial myosin formation and impair apicomedial Rok accumulation in SG cells. (A-B’’’) sqh-GFP signals in control (A-A’’’) and *Rab11* RNAi (B-B’’’) SGs immunostained for E-Cad and GFP. Higher magnification of the yellow boxed area in A and B are shown in A’’-A’’’ and B’’-B’’’. Compared to strong signals of apicomedial myosin web structures in the control SG (yellow arrowheads in A’’), reduced and dispersed myosin signals are detected upon *Rab11* knockdown (red arrowheads in B’’). (A’’’’ and B’’’’) Inverted sqh-GFP signals are used for measuring areas of apicomedial myosin structures. (C-E) Quantification of the intensity of apicomedial (AM) and junctional (J) myosin I, the ratio of apicomedial to junctional myosin (D) and areas of apicomedial myosin (E) in control and *Rab11* RNAi SG cells. n= 5 SGs for both genotypes; 10 cells in the dorsal/posterior region of each SG. *, p< 0.05; **, p< 0.01. Welch’s t-test. (F-G’’) Rok-GFP signals in control (F, F’) and Rab11 RNAi (G, G’) SGs immunostained for E-Cad and GFP. Higher magnification of the yellow boxed area in F and G are shown in F’ and G’. Compared to the strong accumulation of Rok-GFP signals in the apicomedial region of control SG cells (yellow arrowheads in F’), Rok-GFP signals are more dispersed when *Rab11* is knocked down (red arrowheads in G’). (F’’ and G’’) Inverted Rok-GFP signals are used for measuring areas of apicomedial Rok accumulation. (H) Quantification of areas of Rok-GFP puncta. *, p< 0.05; **, p<0.01. Welch’s t-test. n= 5 SGs for all genotypes; 15 cells in the dorsal/posterior region for each SG. Asterisks: invagination pit.

We also tested whether junctional myosin, a myosin pool closely associated with adherens junctions in SG cells, is affected in *Rab11* RNAi SGs. Consistent with previous studies (Roper, 2012; Chung *et al*., 2017), junctional myosin showed strong signals in wild type SG cells, often with stronger signals at vertices (Figure 4A’’’). Intriguingly, the junctional myosin intensity slightly increased when *Rab11* was knocked down (Figure 4, B’’’ and D). The ratio of apicomedial to junctional myosin intensity was significantly reduced in *Rab11* RNAi SGs (Figure 4D), suggesting an imbalance of contractile forces in the SG.

We previously showed that accumulation of apicomedial Rok is required for apicomedial myosin formation for coordinated apical constriction in SG cells (Roper, 2012; Chung *et al*., 2017). To test whether Rok accumulation is dependent on Rab11, we quantified accumulation of apicomedial Rok-GFP signals using a ubiquitously expressed GFP-tagged Rok transgene (Rok-GFP; (Abreu-Blanco *et al*., 2014) in wild type and *Rab11* RNAi SGs (Figure 4, F-G’’). Indeed, the reduction of *Rab11* levels led to more dispersed Rok signals along the apical domain of SG cells near the pit (Figure 4H). Overall, our data suggest Rab11-dependent apical apicomedial myosin formation and accumulation of Rok during SG invagination.

### Reduced dynein function leads to failure in proper organization of apicomedial myosin in SG cells

We next tested whether apicomedial myosin formation is affected in *Dhc64C* RNAi and *klar* mutant SGs. Despite the slight disruption of the MT networks in *Dhc64C* RNAi SGs (Figure 3J’), the overall intensity of apicomedial or junctional myosin was not significantly changed in *Dhc64C* RNAi SGs (Figure 5C), suggesting that subtle defects in the MT networks do not affect apical upregulation of myosin. Similar to *Dhc64C* knockdown, no significant difference in the intensity of apicomedial and junctional myosin was observed in *klar* mutant SGs (Figure 5H). However, the ratio of apicomedial to junctional myosin was reduced in both genotypes (Figure 5, D and I). Moreover, the area of apicomedial myosin accumulation was significantly reduced in *Dhc64C* RNAi or *klar* mutant SGs (Figure 5, E and J), suggesting defects in forming proper apicomedial myosin web in these genotypes. Our data suggest that defective apical constriction in *Dhc64C* RNAi and *klar* mutant SGs could be due to failure in proper organization of myosin structures in SG cells.

**Figure 5.**
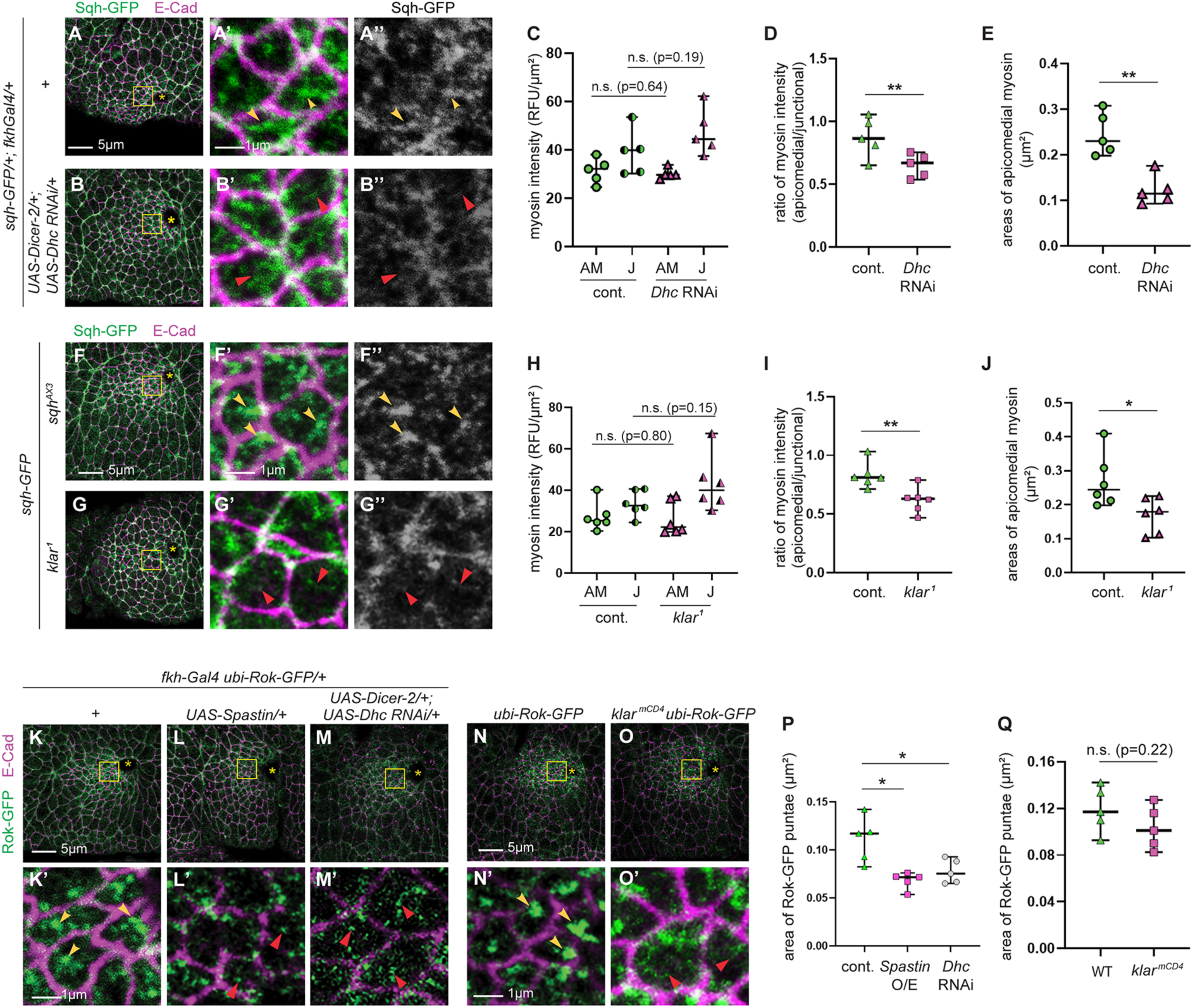
Compromised dynein function leads to a reduction of apicomedial myosin formation and impairs apicomedial Rok accumulation in SG cells. (A-B’’, F-G’’) sqh-GFP signals in control (A-A’’, F-F’’), *Dhc64C* RNAi (B-B’’) and *klar* mutant (G-G’’) SGs immunostained for E-Cad and GFP. Higher magnification of the yellow boxed area in A, B, F and G are shown in A’, A’’, B’, B’’, F’, F’’, G’ and G’. Yellow and red arrowheads indicate apicomedial myosin. (C-E, H-J) Quantification of the intensity of apicomedial and junctional myosin (C, H), the ratio of apicomedial to junctional myosin (D, I) and areas of apicomedial myosin (E, J) in SG cells. n= 5 SGs for all genotypes; 10 cells in the dorsal/posterior region of each SG. *, p< 0.05; **, p< 0.01, Welch’s t-test. (A-A’’) (K-O’) Rok-GFP signals in control (K, K’, N, N’), spastin-overexpressing (L, L’), *Dhc64C* RNAi (M, M’) and *klar* mutant (O, O’) SGs immunostained for E-Cad and GFP. Higher magnification of the yellow boxed area in K-O are shown in K’-O’. (P, Q) Quantification of areas of Rok-GFP puncta. *, p< 0.05; **, P<0.01, Welch’s t-test. n= 5 SGs for all genotypes; 15 cells in the dorsal/posterior region for each SG. Asterisks: invagination pit.

We also tested for accumulation of apicomedial Rok-GFP signals in spastin-overexpressing, *Dhc64C* RNAi and *klar* mutant SGs. Disruption of MTs by spastin overexpression abolished apicomedial accumulation of Rok in cells near the invagination pit (Figure 5, L’ and P), found in control SGs (Figure 5K’). Reduction of *Dhc64C* levels also led to more dispersed Rok signals along the apical domain of SG cells near the pit (Figure 5, M’ and P). In *klar* mutants, Rok-GFP tended to accumulate less profoundly and less uniformly in the apicomedial region of SG cells (Figure 5O’) although not statistically significant compared to control (Figure 5Q). Overall, our data suggest that MT- and dynein-dependent apical accumulation of Rok and apicomedial myosin formation during SG invagination.

### MT- and Rab11-dependent apical enrichment of Fog, Crb and E-Cad is required for apical constriction during SG invagination

We next tested potential cargos of MT- and Rab11-dependent apical transport during SG invagination. Rok accumulation, apicomedial myosin formation and subsequent clustered apical constriction in the SG is dependent on Fog signaling (Chung *et al*., 2017). We therefore tested if apical transport of the Fog ligand is MT- and Rab11-dependent. Staining using an antibody against Fog in control SGs showed upregulated Fog signals in the apical domain of SG cells (Figure 6, A-A’’’). However, in spastin-overexpressing or *Rab11* RNAi SGs, apical Fog signals were reduced (Figure 6, B-C’’’). To compare apical enrichment of Fog between WT and spastin-overexpressing or *Rab11* RNAi SGs, we calculated the overall intensity of apical Fog signals in the entire SG placode in corresponding genotypes. Compared to control, Fog levels in the apical domain of the SG were reduced when spastin was overexpressed or *Rab11* was knocked down (Figure 6D). We also calculated the degree of variability, the ratio of the deviation of Fog intensity to the mean intensity of Fog in the SG. Compared to WT, spastin-overexpressing or *Rab11 RNAi* SGs had a lower degree of relative variability of intensity (Figure 6E). These data suggest that apical Fog signals are less varied from cell to cell regardless of apical areas when MTs are disrupted or Rab11 function is reduced. Overall, these data suggest that apical transport of Fog in the SG is dependent on the MT networks and Rab11.

**Figure 6.**
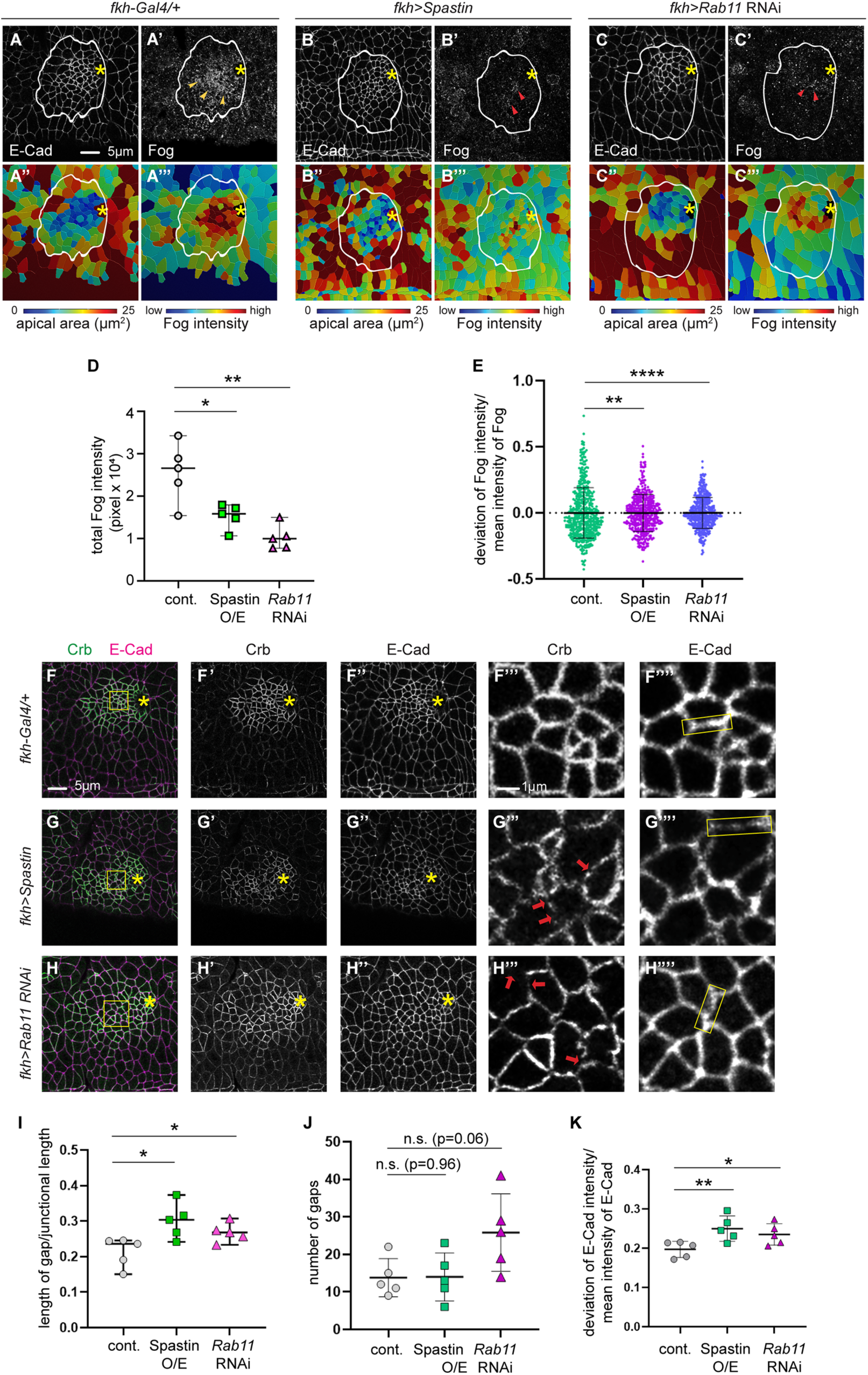
MTs and Rab11 are required for apical transport of Fog, Crb and E-Cad during SG invagination. (A-C’) Confocal images for control (A-A’), spastin-overexpressing (B-B’) and *Rab11* RNAi (C-C’) SGs immunostained for E-Cad and Fog. (A’’-C’’’) Corresponding heat maps for apical areas (A’’-C’’) and Fog intensity (A’’’-C’’’). (D) Total intensity of apical Fog signals in the entire SG placode in control, spastin-overexpressing and *Rab11* knocked down SG. n= 5 SGs for each genotype. (E) The degree of variability for Fog signals. n= 5 SGs (control (*fkh-Gal4/+*), 475 cells; *fkh-Gal4/UAS-spastin*, 507 cells; *fkh-Gal4/UAS-Rab11* RNAi, 487 cells). Kolmogorov-Smirnov test. (F-H’’’’) Confocal images for control (F-F’’’’), spastin-overexpressing (G-G’’’’) and *Rab11* RNAi (H-H’’’’) SGs immunostained for E-Cad and Crb. Higher magnification of boxed areas in F-H are shown in F’’’-H’’’’. Compared to relatively continuous Crb signals in the control SG (F’’’), Crb signals show gaps (red arrows) in spastin-overexpressing (G’’’) and *Rab11* RNAi (H’’’) SGs. Compared to relatively uniform E-Cad signals along adherens junctions in the control SG (F’’’’), E-Cad signals in spastin-overexpressing (G’’’’) and *Rab11* RNAi (H’’’’) SGs are unevenly distributed. Yellow boxes, representative junctions. (I) Quantification of the ratio of length of gaps to junctional length (Welch’s t-test) in SGs immunostained for Crb. (J) Quantification of the number of gaps (Welch’s t-test) in SGs immunostained for Crb. n= 5 SGs; 10 cells for each SG. (K) The degree of variability of E-Cad intensities in the SG. n= 5 SGs; 20 junctions for each SG. Welch’s t-test. *, p<0.05; **, p< 0.01; ****, p< 0.0001. Asterisks: invagination pit. White lines: SG boundary.

We further tested MT- and Rab11-dependent apical transport of other apical and junctional proteins during SG invagination. Transport of several key apical and junctional proteins is dependent on MTs and Rab11 (Le Droguen *et al*., 2015; Khanal *et al*., 2016; Jouette *et al*., 2019). One such protein is an apical transmembrane protein Crb, and Rab11 helps maintain apical Crb in the *Drosophila* ectoderm (Roeth *et al*., 2009). In the *Drosophila* SG and follicle cells, Crb is apically transported along MTs by the dynein motor (Myat and Andrew, 2002). Importantly, in *Drosophila* tracheae, loss of *crb* impairs apical constriction during the internalization process (Letizia *et al*., 2011). The key junctional protein E-Cad is also trafficked by MTs in *Drosophila* embryonic trachea (Le Droguen *et al*., 2015). During apical constriction, contractile forces generated by the actomyosin complex are exerted on adherens junctions, with E-Cad being a core component that integrates contractile forces to generate tension (Martin *et al*., 2009).

We therefore asked if MTs and Rab11 have roles in apical distribution of Crb and E-Cad in the SG during invagination. Compared to control (Figure 6, F’ and F’’’), spastin-overexpressing SG cells showed discontinuous Crb signals (Figure 6, G’ and G’’’). Although the number of gaps was not significantly different between control and spastin-overexpressing SGs (Figure 6J), the ratio of the length of gap over junctional length was significantly increased in SGs that overexpress spastin (Figure 6I). Unlike for Crb, disruption of MTs in the SG did not cause obvious gaps in E-Cad signals at adherens junctions compared to control (Figure 6, F’’, F’’’’, G’’ and G’’’’). However, we observed an ununiform distribution of E-Cad signals along adherens junctions in spastin-overexpressing SGs (Figure 6G’’’’). The degree of variability of E-Cad intensity along the junctions was significantly increased in spastin-overexpressing SGs compared to control (Figure 6K). Importantly, *Rab11* knockdown resulted in similar discontinuous Crb and uneven E-Cad signals (Figure 6, H-H’’’’, I and K). These data suggest that proper apical distribution of Crb and E-Cad in the invaginating SG is dependent on MTs and Rab11.

To test for roles of Crb and E-Cad in regulating apical constriction during SG invagination, we knocked down either gene in the SG using RNAi. Knockdown of *crb* or *E-Cad* resulted in the reduction of junctional Crb or E-Cad levels, respectively, compared to control (Supplemental Figure S3, E-F’’). Reduction of *crb* increased the number of cells with larger apical areas compared to control (Figure 7, A, B and D). Quantification of the percentage of cells and cumulative percentage of cells of different apical areas showed a significant decrease in the number of constricting cells in *crb* knockdown SGs (Figure 7D). *E-Cad* knockdown SGs also displayed a similar trend, although with less statistical significance (Figure 7, C and D). These data suggest roles for Crb and E-Cad, with more subtlety for E-Cad, in regulating apical constriction during SG invagination.

**Figure 7.**
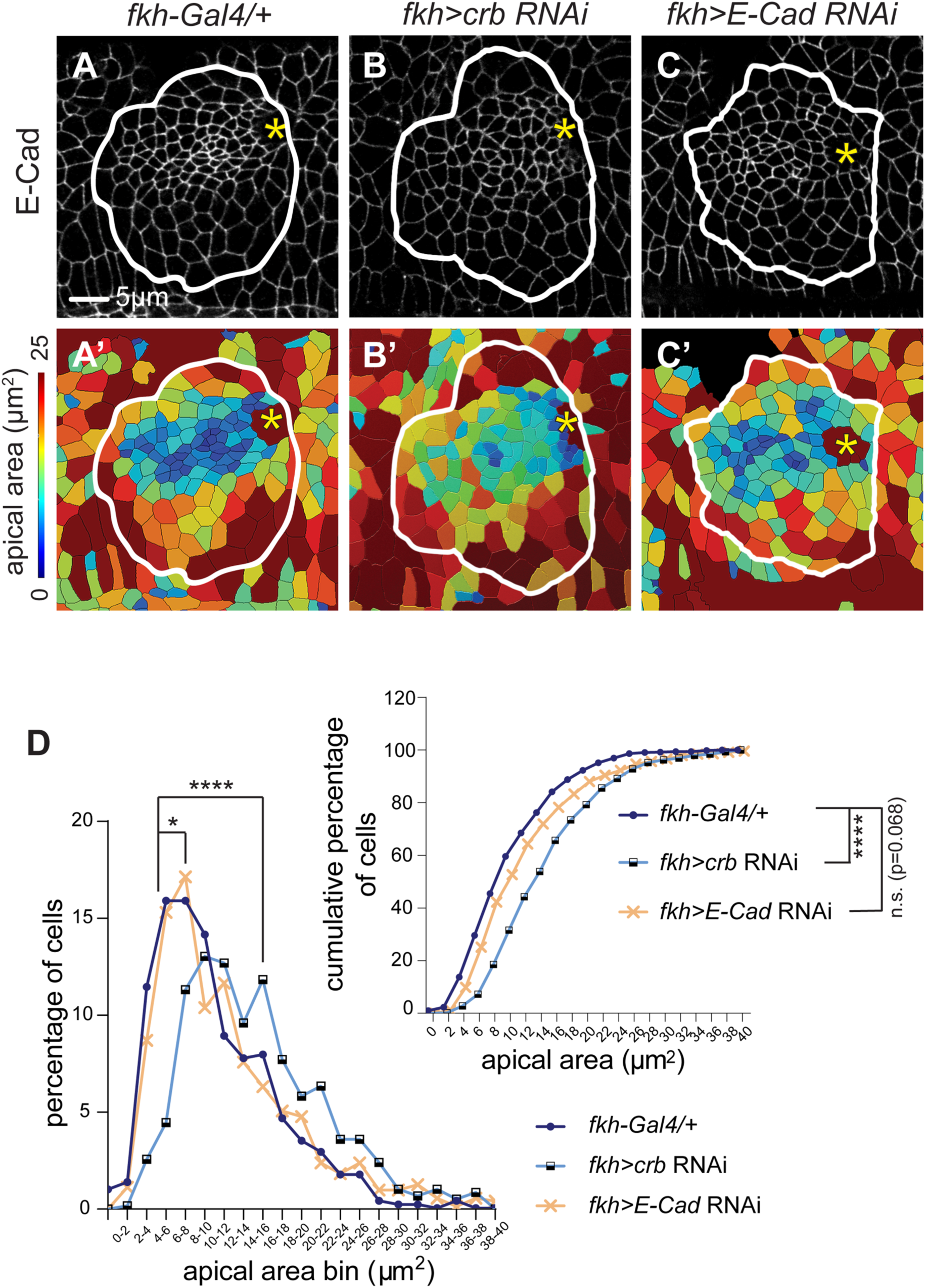
Crb and E-Cad have roles in regulating apical constriction during SG invagination. (A-C) Confocal images of control (A), *crb* RNAi (B) and *E-Cad* RNAi (C) SGs immunostained for E-Cad. (A’-C’) Corresponding heat maps for apical areas in SGs in A-C. (D) Percentage and cumulative percentage of cells with different apical areas. Mann-Whitney U test (percentage of cells) and Kolmogorov-Smirnov test (cumulative percentage of cells). n= 5 SGs (control, 517 cells; *crb* RNAi, 583 cells; *E-Cad* RNAi, 712 cells). *, p<0.05; ****, p<0.0001. Asterisks: invagination pit. White lines: SG boundary.

### Apical localization of Fog, Crb and E-Cad is MT- and Rab11-dependent in the SG throughout development

To further investigate the role of MTs in Rab11-dependent apical trafficking during SG development, we overexpressed spastin in the SG throughout development and analyzed late stage SGs. Analysis of spastin-overexpressing SGs at stage 16 revealed that disruption of MTs resulted in short SG tubes with a thin lumen compared to control SGs of the same stage (Supplemental Figure S4, A and B). Strikingly, whereas the majority of Rab11- and Nuf-positive vesicles localized in the apical region of control SG cells (Supplemental Figure S4, A-A’’’), Rab11 and Nuf were observed as large aggregates in the cytoplasm in spastin-overexpressing cells, overlapping with each other (Supplemental Figure S4, B-B’’’). Our data suggest that MTs are required for apical enrichment of Rab11/Nuf vesicles throughout SG tube formation.

We next tested whether MTs and Rab11 are required for apical transport of Fog, Crb and E-Cad also at later stages. Compared to the control SG (Supplemental Figure S4C’’), apical Fog signals were significantly reduced in SGs that overexpress spastin or knock down *Rab11* (Supplemental Figure S4, D’’ and E’’), suggesting that Fog is trafficked apically in a MT- and Rab11-dependent manner in the SG throughout tube formation. No significant changes were detected in apical Crb localization in SGs that overexpress spastin or knock down *Rab11* (data not shown). Compared to strong E-Cad enrichment in adherens junctions in control SG cells (Supplemental Figure S4C’), SGs that overexpress spastin or knock down *Rab11* showed stronger E-Cad signals in the lateral domain of cells (Supplemental Figure S4, D’ and E’).

To further test MT-dependent apical transport of Crb or E-Cad in the SG, we used a genetic suppression strategy in the overexpression background: we co-overexpressed Crb or E-Cad along with spastin in the SG to test whether disruption of MTs affects apical transport of Crb or E-Cad. A similar approach was taken in a recent study, where overexpression of Crb provided a highly sensitive genetic background for identifying components involved in Crb trafficking (Aguilar-Aragon *et al*., 2020). Consistent with previous studies (Wodarz *et al*., 1993; Chung and Andrew, 2014), Crb overexpression in the SG caused a dramatic increase of the apical membrane as well as mislocalization of Crb basolaterally along the entire membrane (Supplemental Figure S4, F-F’). Co-overexpression of spastin along with Crb suppressed the overexpression phenotypes for Crb (Supplemental Figure S4, G-G’). E-Cad overexpression in the SG caused mislocalization of E-Cad as punctate structures in the basolateral region (Supplemental Figure S4, H’-H’’). Importantly, in SGs that co-overexpressed spastin along with Crb or E-Cad, each protein mislocalized as cytoplasmic aggregates near the basolateral domain, which were largely overlapping with mislocalized Rab11/Nuf (Supplemental Figure S4, G’-G’’’ and I’-I’’’). Overall, our data suggest that Fog, Crb and E-Cad are trafficked apically in a MT- and Rab11-dependent manner in the SG throughout tube formation.

### Reducing apical and junctional proteins affects apicomedial myosin formation

To test whether apical constriction defects observed in *crb* and *E-Cad* RNAi SGs are linked to apicomedial myosin formation, we quantified myosin levels and areas of apicomedial myosin in SGs with these knockdowns. Compared to control SGs (Figure 8, A-A’’), *crb* RNAi SGs showed a slight decrease in apicomedial myosin levels (Figure 8, B-B’’ and D). No significant difference in the junctional myosin intensity was observed in *crb* RNAi SGs (Figure 8, B-B’’ and D). The ratio of apicomedial to junctional myosin was reduced (Figure 8E), suggesting that apical constriction defects in SGs depleted of *crb* are, at least in part, due to failure of apicomedial myosin formation. Supporting this idea, areas of apicomedial myosin web-like structures were significantly reduced in *crb* RNAi SGs compared to WT (Figure 8F).

**Figure 8.**
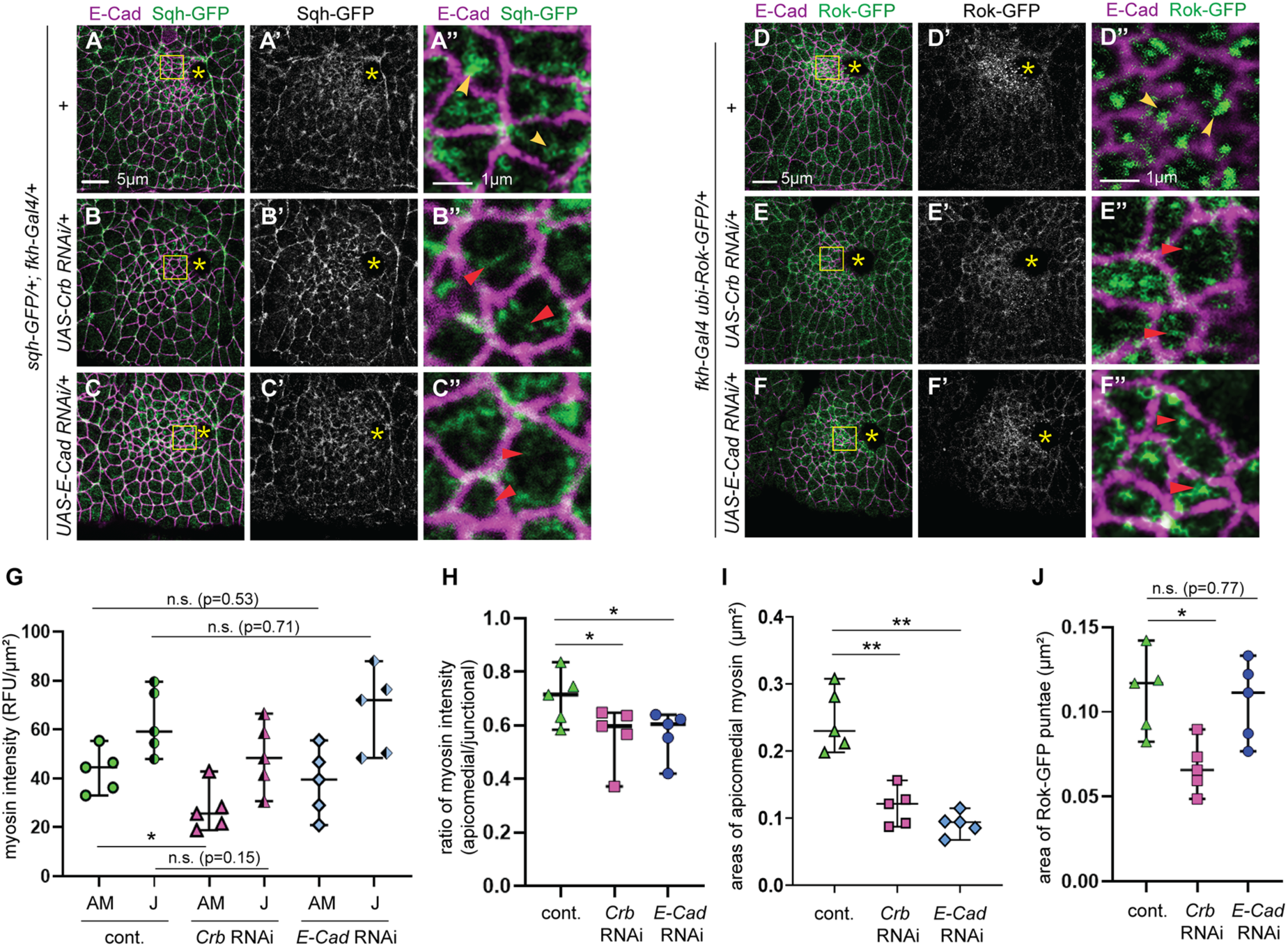
Knockdown of *crb* or *E-Cad* results in reduced apicomedial myosin formation and dispersed apicomedial Rok. (A-C”) sqh-GFP signals in control (A-A’’), *crb* RNAi (B-B’’) and *E-Cad* RNAi (C-C’’) SGs immunostained for E-Cad and GFP. (A”- C”) Higher magnification of yellow boxed areas in A-C. (D-F”) Rok-GFP signals in control (D-D’’), *crb* RNAi (E-E’’) and *E-Cad* RNAi (F-F’’) SGs immunostained for E-Cad and GFP. (D”-F”) Higher magnification of yellow boxed areas in D-F. (G-I) Quantification of the intensity of apicomedial and junctional myosin (G), the ratio of apicomedial to junctional myosin (H) and areas of apicomedial myosin structures (I) in SG cells from different genotypes shown in A-C (n=5 SGs for each genotype; 10 cells in the dorsal/ posterior region). *, p<0.05; **, p<0.01. Welch’s t-test. (F-I’’) (J) Quantification of areas of Rok-GFP puncta in SG cells from different genotypes shown in D-F (n=5 SGs; 15 cells in the dorsal/posterior for each genotype). *, p<0.05; **, p< 0.01. Welch’s t-test. Asterisks: invagination pit.

**Figure 9.**
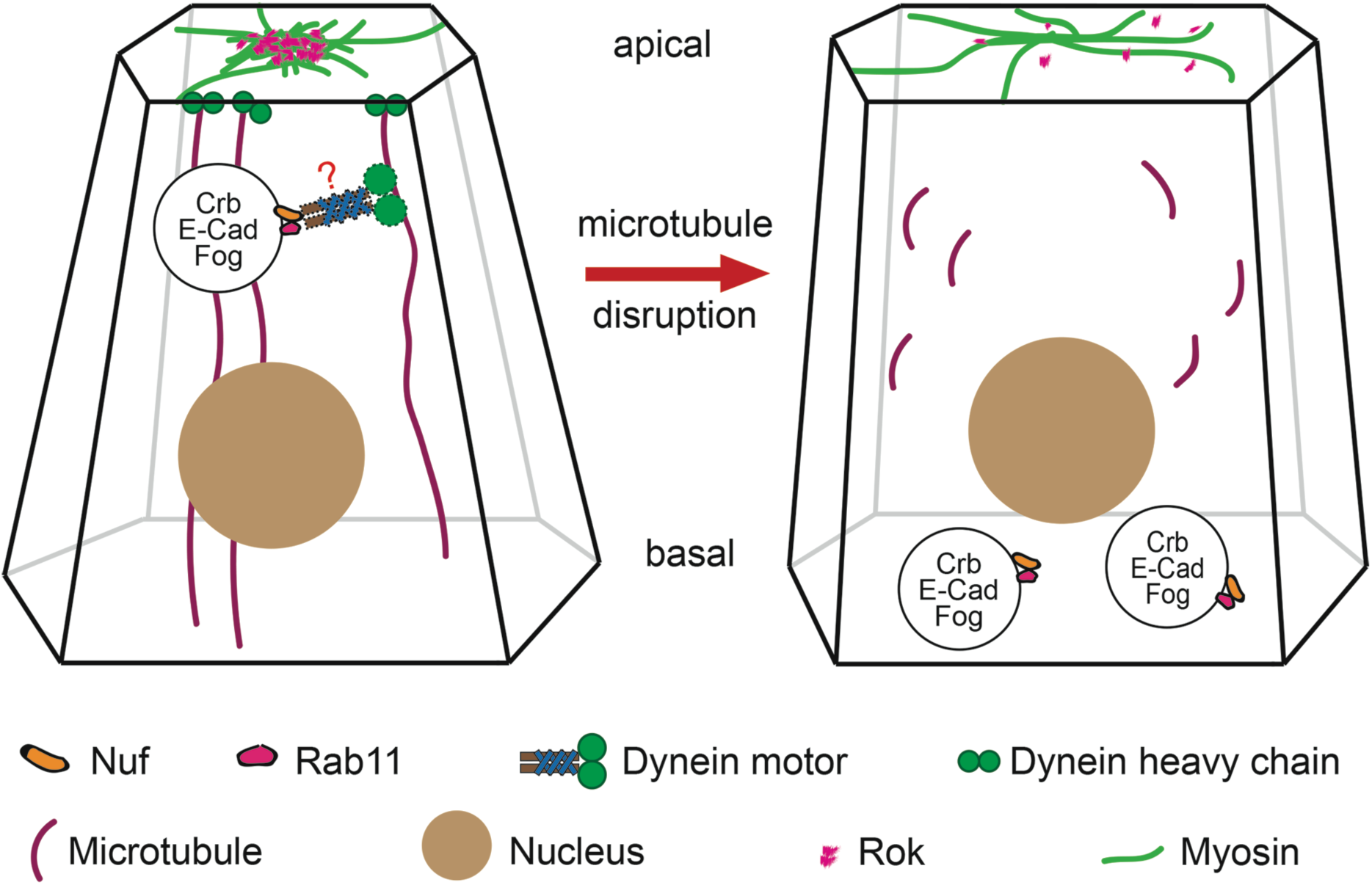
A proposed model for MT-dependent trafficking to promote apical constriction during SG invagination. Vesicular transport is essential for apical localization of Fog, Crb and E-Cad to regulate apical myosin networks and subsequent apical constriction. Precise roles for the dynein motor in regulating apical constriction remain to be determined. The red question mark represents a potential role of dynein in trafficking cargos to the apical region of SG cells during apical constriction.

In SGs knocked down *E-Cad*, myosin levels at each domain did not change significantly (Figure 8, C-C’’ and E), but areas of apicomedial myosin web-like structures were significantly decreased (Figure 8F). The ratio of apicomedial to junctional myosin was also significantly reduced in *E-Cad* RNAi SGs compared to control (Figure 8, C-C’’ and E). Therefore, apical constriction defects caused by *E-Cad* knockdown might be due to defective apicomedial myosin formation and the imbalance of contractile forces.

Consistent with reduced apicomedial myosin (Figure 8, B’ and D), knockdown of *crb* also resulted in Rok-GFP signals less accumulating in the apical region of cells near the invagination pit (Figure 8, F-G’). Quantification of areas occupied by Rok-GFP puncta showed significant reduction in accumulation of apicomedial Rok when *crb* was knocked down (Figure 8I). Consistent with little effect of *E-Cad* knockdown on apicomedial myosin intensity (Figure 8, C-C’’ and E), Rok accumulation was not significantly affected by *E- Cad* knockdown (Figure 8, H-H’ and I). Taken together, our results suggest a role for Crb in regulating apical constriction by apicomedial myosin activation during SG invagination.

## Discussion

### MTs and Rab11 regulate Fog signaling activity during SG invagination

MTs have a crucial role in stabilizing apical myosin during epithelial morphogenesis both in early *Drosophila* embryos and in the *Drosophila* SG (Booth *et al*., 2014; Ko *et al*., 2019). In the SG, MTs interact with apicomedial myosin via Short stop, the *Drosophila* spectraplakin, emphasizing a direct interplay between the MT and the apical myosin networks (Booth *et al*., 2014). Our data reveals another key role of MTs in regulating protein trafficking to control the apical myosin networks during tissue invagination. During SG invagination, a network of longitudinal MT bundles is observed near the invagination pit (Booth *et al*., 2014). Our data shows apical enrichment of Rab11 in the same area is MT-dependent (Figures 1 and 2) and that this enrichment is important for forming the apicomedial myosin networks (Figure 4), suggesting a link between localized intracellular trafficking along MTs to apical myosin regulation.

The dorsal/posterior region of the SG, where Rab11 is apically enriched in a MT-dependent manner (Figures 1 and 2), correlates with localized Fog signaling activity that promotes clustered apical constriction (Chung *et al*., 2017). Disruption of MTs or *Rab11* knockdown reduces Fog signals in the apical domain of SG cells (Figure 6) and causes dispersed Rok accumulation and defective apicomedial myosin formation (Figure 4). It is consistent with our previous study that the absence of Fog signal results in dispersed apical Rok and defects in apicomedial myosin formation (Chung *et al*., 2017). We therefore propose that MT- and Rab11-dependent apical trafficking regulates Fog signaling activity to control apical constriction during epithelial tube formation, through transporting the Fog ligand. As recycling of membrane receptors to the cell surface plays an important role in regulating overall signaling activity, it is possible that Rab11 is involved in recycling the as yet unidentified SG receptor(s) of Fog to regulate Fog activity in the SG. Indeed, several GPCRs are recycled via Rab11 (Anborgh *et al*., 2000; Innamorati *et al*., 2001; Hunyady *et al*., 2002; Volpicelli *et al*., 2002; Fan *et al*., 2003; Dale *et al*., 2004; Hamelin *et al*., 2005; Cerniello *et al*., 2017). During epithelial invagination in early *Drosophila* embryogenesis, the concentration of the Fog ligand and receptor endocytosis by *β*-arrestin-2 have been shown as coupled processes to set the amplitude of apical Rho1 and myosin activation (Jha *et al*., 2018). It is possible that the movement of Fog receptor(s) that have internalized as a stable complex with *β*-arrestin is recycled back to the cell surface by Rab11. The Fog signaling pathway represents one of the best-understood signaling cascades controlling epithelial morphogenesis (Manning and Rogers, 2014). Although best studied in *Drosophila*, the pathway components have also been identified in other insects, suggesting a more widely conserved role of Fog signaling in development (Urbansky *et al*., 2016; Benton *et al*., 2019). Further work needs to be done to fully understand the regulatory mechanisms underlying the trafficking of Fog and its receptor(s) during epithelial morphogenesis.

Our analysis of apicomedial myosin shows that reduced Rab11 function not only causes a decrease of the myosin intensity but also causes myosin to be dispersed rather than forming proper myosin web structures in the apicomedial domain of SG cells (Figure 4). These data support the idea that Rab11 function is required for both concentration and spatial organization of apicomedial myosin. This can be explained by the combined effect of multiple cargos that are transported by Rab11, including Fog, Crb and E-Cad (as discussed below). Time-lapse imaging of myosin will help determine how the dynamic behavior of apicomedial myosin is compromised when Rab11 function is disrupted.

### Integrating apical and junctional proteins with actomyosin networks during SG invagination

During branching morphogenesis in *Drosophila* trachea, MTs and dynein motors have a critical role in the proper localization of junctional proteins such as E-Cad (Le Droguen *et al*., 2015). This is consistent with our observations with MT-dependent uniform distribution of E-Cad at adherens junctions in the invaginating SG (Figure 6), suggesting a conserved role of MT-dependent intracellular trafficking in junctional remodeling and stabilization during epithelial tube formation. Our data further suggest that the MT networks and Rab11 have key roles in apical distribution of Crb and E-Cad in the SG (Figure 6) and that proper levels of apical and junctional proteins are important for apical constriction during SG invagination (Figure 7). Based on these data, we propose that MT- and Rab11-dependent apical trafficking of Crb and E-Cad is critical for apical constriction during SG invagination. Alternatively, MTs have an additional role in assembling/anchoring these apical components, through the regulation of unidentified molecules. Recent studies in *Drosophila* mesoderm invagination showed that MTs help establish actomyosin networks linked to cell junction to facilitate efficient force transmission to promote apical constriction (Ko *et al*., 2019). In (Ko *et al*., 2019), however, MT-interfering drugs and RNAi of CAMSAP end-binding protein were used to prevent MT functions and the effect cannot be directly compared to our data where spastin was used to sever existing MTs. Direct monitoring of MT-dependent transport of Crb and E-Cad during SG invagination will help clarify the mechanism.

Upon knockdown of *crb* or *E-Cad,* less prominent apicomedial myosin web structures are observed in invaginating SGs (Figure 8), suggesting a requirement of Crb and E-Cad in proper organization of apical actomyosin networks during SG tube formation. Crb acts as a negative regulator of actomyosin dynamics during *Drosophila* dorsal closure (Flores-Benitez and Knust, 2015) and during SG invagination (Roper, 2012). It is possible that proper Crb levels are required for modulating myosin activity both in the apicomedial domain and at junctions during SG invagination, which contribute to apical constriction and cell rearrangement, respectively (Roper, 2012; Sanchez-Corrales *et al*., 2018). Anisotropic localization of Crb and myosin was observed at the SG placode boundary, where myosin accumulates at edges where Crb is lowest (Roper, 2012). Planar polarization of Rok at this boundary is modulated through phosphorylation by Pak1 downstream of Crb (Sidor *et al*., 2020). A further test will help understand whether and how Crb might affect junctional myosin dynamics and SG invagination. As contractile actomyosin structures exert forces on adherens junction to drive apical constriction, we speculate that apical constriction defects upon *E-Cad* RNAi might be due to reduction of cell adhesion and/or of improper force transmission. It will be interesting to determine if the coordination of apical and junctional proteins and apical cytoskeletal networks through intracellular trafficking is conserved during tubular organ formation in general.

### A role for dynein in apical constriction

Dhc is also apically enriched in the dorsal/posterior region of the invaginating SG (Figure 1). Our data show that knockdown of *Dhc64C* not only affects Rok accumulation and apicomedial myosin formation (Figure 5) but also disrupts MT organization in the SG (Figure 3). This data is consistent with previous findings that cytoplasmic dynein is associated with cellular structures and exerts tension on MTs. For example, dynein tethered at the cell cortex can apply a pulling force on the MT network by walking towards the minus end of a MT (Laan *et al*., 2012). Dynein also scaffolds the apical cell cortex to MTs to generate the forces that shape the tissue into a dome-like structure (Takeda *et al*., 2018). In interphase cells, the force generated by dynein also regulates MT turnover and organization (Yvon *et al*., 2001).

In *klar* mutants, on the other hand, MT organization is not affected in the SG (Figure 3), suggesting that reduction of dynein-dependent trafficking by loss of *klar* does not cause changes in the MT networks. Notably, although the intensity of apicomedial myosin does not change upon *Dhc64C* knockdown or in the *klar* mutant background, formation of apicomedial myosin web structures is affected (Figure 5). These data suggest a possible scenario that dynein function is not required for myosin concentration in the apical domain but is only needed for the spatial organization of apicomedial myosin. However, we cannot rule out the possibility that the zygotic knockdown of *Dhc64C* by RNAi is not strong enough to affect the intensity of apicomedial myosin. *Dhc64C* has strong maternal expression and is essential for oogenesis and early embryo development (Li *et al*., 1994). Embryos with reduced maternal and zygotic pools of *Dhc64C* showed a range of morphological defects in the entire embryo, some of which were severely distorted (data not shown). Precise roles for dynein and dynein-dependent trafficking in regulating apicomedial myosin formation remain to be elucidated.

## Materials and Methods

### Fly stocks and husbandry

Fly lines used in our experiments were listed in a separate table. All crosses were performed at 25°C, unless stated otherwise.

### Antibody staining and confocal microscopy

Antibodies used in our experiments were listed in a separate table. Embryos were collected on grape juice-agar plates and processed for immunofluorescence using standard procedures. Briefly, embryos were dechorionated in 50% bleach, fixed in 1:1 heptane:formaldehyde for 40 min and devitellinized with 80% EtOH, then stained with primary and secondary antibodies in PBSTB (1X PBS, 0.1% Triton X-100, 0.2% BSA). For acetylated α-tubulin, tyrosinated α-tubulin, sqh-GFP, Rok-GFP, Fog and phalloidin staining, embryos were hand-devitellinized. All images were taken with a Leica SP8 confocal microscope.

### Cell segmentation and apical area quantification

Embryos immunostained with E-Cad and CrebA were imaged using a Leica SP8 confocal microscope. As Rok accumulation, apicomedial myosin and apical constriction depend on the depth of invagination in the SG, SGs that were invaginated within the range of 5.1-9.9 μm depth were used for quantification for proper comparison between different genotypes. Maximum intensity projection was generated from three apical focal planes with the highest E-Cad signals for all genotypes (0.3 μm apart for each focal plane). Cells were segmented along E-Cad signals and cell areas were calculated using the Imaris Program (Bitplane). Since the Imaris Program calculated the areas of both the front and the back of the projected cell layer, we divided the measured areas by two to get the true values of apical areas of SG cells.

### Negative correlation between apical area and Rab11/Nuf intensity

Cell segmentation for five WT SGs within the range of 5.1-9.9 μm invagination was performed as described above. All experiments were carried out in the same condition, and the same settings for confocal imaging were used. Three confocal sections in the apical region that show the strongest Rab11/Nuf signals were used to produce the maximum intensity projection. Intensity means were measured for Rab11/Nuf signals for each segmented cell using the Imaris Program (Bitplane) and plotted. Correlation (Pearson) and P values were calculated using the GraphPad Prism software.

### Total intensity of Rab11, Nuf and Fog signals

For measuring the total intensity of Rab11/Nuf signals in the apical region of cells in the whole SG placode, the integrated density of Rab11/Nuf of each SG cell was calculated (integrated density= intensity mean x area). Intensity means and SG cell areas were measured using the Imaris software. The total intensity was calculated as the sum of integrated densities of all cells in the placode. Five SGs were used for quantification. P values were calculated using Welch’s t-test in the GraphPad Prism software.

For the total intensity of Fog signals, background corrections were carried out first. Using the Fiji software, mean gray values of Fog signals of ten epidermal cells outside of the SG placode were measured. The background intensity was calculated as the average value of the intensity means of those ten cells. The intensity mean of Fog signals of all SG cells was measured using the Imaris program, and the background intensity was subtracted. After background correction, the intensity mean was multiplied by SG cell areas to calculate the total Fog intensity. Five SGs were used for quantification. P values were calculated using Welch’s t-test in the GraphPad Prism software.

### The degree of variability for Rab11, Nuf, Fog and E-Cad signals

The degree of variability of Rab11/Nuf signals was calculated as the ratio of deviation of Rab11/Nuf intensity to the mean Rab11/Nuf intensity. The mean value of Rab11/Nuf intensity was calculated as the average mean intensity of Rab11/Nuf signals in all SG cells. Deviation of Rab11/Nuf intensity is the difference between the mean intensity of Rab11/Nuf in each cell and the mean value. Five WT SGs (690 cells) and five spastin-overexpressed SGs (496 cells) were analyzed and plotted. P values were calculated using the Mann-Whitney U test in the GraphPad Prism software.

The degree of variability of Fog signals in control, spastin-overexpressing and Rab11 RNAi SGs was calculated using the same methods for Rab11/Nuf. Five SGs for each genotype were analyzed (control (*fkh-Gal4/+*), 475 cells; *fkh-Gal4/UAS-Spastin*, 507 cells; *fkh-Gal4/UAS-Rab11* RNAi, 487 cells).

The degree of variability for E-Cad signals was calculated as the ratio of average deviation of E-Cad to the mean E-Cad intensity. To measure the intensity of E-Cad signals along the adherens junction, we drew a polyline along E-Cad signals at each junction. The deviation and mean intensity of E-Cad signals were measured using the Leica LasX software. SuperPlots (Lord *et al*., 2020) were used to display the quantification. Each dot in the graph represents the average value of 20 junctions in the dorsal posterior region of each SG. Five SGs were analyzed for each genotype. P values were calculated using Welch’s t-test in the GraphPad Prism software.

### Quantification of intensities of myosin, Crb and E-Cad

For myosin quantification, maximum intensity projections that span the apical and the junctional region of SG cells were used (Leica LasX) and measurements were performed using the Fiji software. Five SGs were used for quantification for each genotype. A group of 20 cells in the dorsal/posterior region of the SG placode near the invagination pit was selected for quantification of myosin intensity. Regions were drawn manually along the inner or outer boundary of E-Cad signals of each cell to calculate the mean gray value of apicomedial and junctional myosin.

For background correction, mean gray values of apicomedial myosin in ten cells outside of the SG placode were measured. The average value of mean gray values of apicomedial myosin in these ten cells was used to subtract the background of the cells inside the placode from the same embryo. The mean intensity of apicomedial/junctional myosin was normalized by the median deviation. SuperPlots (Lord *et al*., 2020) were used to display quantification. For myosin intensity, each dot in the graph represents the average value of 20 cells in each SG. For Crb and E-Cad intensities, each dot in the graph represents the average value of 10 cells for each SG. Five SGs were used for the quantification of all three proteins. P values were calculated using Welch’s t-test in the GraphPad Prism software.

### Quantification of areas of Rok-GFP and apicomedial myosin puncta

For quantification of area of Rok-GFP puncta, a single confocal section that had the strongest medial Rok signals was selected. Cell boundaries were labeled by immunostaining with the antibody against E-Cad. To analyze Rok distribution, we performed Particle Analysis using the Fiji software. Fifteen cells in the dorsal/posterior region near the invagination pit were selected for quantification. Rok-GFP signals were converted into black-and-white using the *Threshold* tool in Photoshop before analysis. Using the *Analyze particles* tool in Fiji, Rok-GFP puncta with areas equal or larger than 0.02 μm^2^ were measured. Five SGs were used for quantification. P values were calculated using Welch’s t-test in the GraphPad Prism software.

For measuring areas of apicomedial myosin puncta, a single confocal section with the strongest apicomedial myosin signals was selected. A group of 10 cells in the dorsal/posterior region of SG was analyzed. Junctional myosin was excluded manually and only apicomedial myosin signals were used for quantification. Measuring the area of apicomedial myosin were carried out with the same method used for quantifying the area of Rok-GFP puncta. Five SGs were used for quantification. P values were calculated using Welch’s t-test in the GraphPad Prism software.

### Quantification of length and number of gaps of junctional Crb

Length of gaps and junctional length were measured using the Fiji software. Gaps that have a length equal or more than 0.2 μm were quantified. If there were more than one gap per junction, the length of gaps was calculated as a sum of all gaps in a given junction. For each SG, ten cells in the dorsal/posterior region were used for quantification. For the number of gaps, the total number of gaps in those ten cells was counted. Five SGs (∼110 junctions) were used for quantification. P values were calculated using Welch’s t-test in the GraphPad Prism software.

**Table 1.** List of fly lines used in this study.

| **Fly strains** | **Source** | **RRID** | **References** |
| --- | --- | --- | --- |
| *Rab11-EYFP* | Bloomington Stock Center | 62549 | (Dunst *et al.*, 2015) |
| *Rab5-EYFP* | Bloomington Stock Center | 62543 | (Dunst *et al.*, 2015) |
| *Rab7-EYFP* | Bloomington Stock Center | 62545 | (Dunst *et al.*, 2015) |
| *UAS-Spastin* (on II) | N. Sherwood, Duke |  | (Sherwood *et al.*, 2004) |
| *UAS-Spastin-CFP* (on III) | N. Sherwood, Duke |  | (Du *et al.*, 2010) |
| *fkh-Gal4* (on II) | D. Andrew (John Hopkins University) |  | (Chung *et al.*, 2017) |
| *fkh-Gal4* (on III) | D. Andrew (John Hopkins University) |  | (Chung *et al.*, 2017) |
| *ubi-Rok-GFP* | S. Parkhurst (Fred Hutchinson Cancer Research Center) |  | (Abreu-Blanco *et al.*, 2014) |
| *sqh-GFP* |  |  | (Royou *et al.*, 2004) |
| *UAS-Rab11S25N-EYFP* | Bloomington Stock Center | 9792, 23261 |  |
| *UAS-Dicer-2* | Bloomington Stock Center | 60008 |  |
| *UAS-Dhc64C* RNAi | Bloomington Stock Center<br>Vienna *Drosophila* Resource Center | 76941, 28749, v28053, v28054.<br>4 lines worked equally.<br>V28054 was used for all of the data shown in this work. |  |
| *Klar*<sup>1</sup> | Bloomington Stock Center | 3256 |  |
| *klar*<sup>mCD4</sup> | Bloomington Stock Center | 25097 |  |
| *klar*<sup>mCD4</sup> *ubi-Rok-GFP* | Recombinant line generated from *klar*<sup>mCD4</sup> and *ubi-Rok-GFP* |  |  |
| *UAS-Crb* RNAi | Bloomington Stock Center | 38373, 38903, 40869, 34999. |  |
|  |  | 4 lines worked equally. 2 lines (38373, 34999) were used for the data shown in this work. |  |
| *UAS-E-Cad*<br>RNAi | Bloomington Stock Center | 32904 |  |
| *UAS-Shg-GFP* | Bloomington Stock Center | 58445 |  |
| *UAS-Rab11*<br>RNAi | Bloomington Stock Center | 42709, 27730.<br>Line 42709 was used for testing myosin and Rok-GFP signals. Line 27730 was used for the rest of the experiments. |  |

**Table 2.** List of primary and secondary antibodies used in this study.

| **Antibody** | **Source** | **RRID** | **Dilution** |
| --- | --- | --- | --- |
| α-E-Cad (rat) | DSHB (deposited by T. Uemura, Kyoto University) | DCAD2<br>(AB_528120) | 1:50 |
| α-CrebA (rat) | Andrew lab (John Hopkins University) | | 1:3000 |
| α-CrebA (rabbit) | Andrew lab (John Hopkins University) | | 1:5000 |
| α-Dhc (mouse) | DSHB (deposited by J.M. Scholey, University of California) | 2C11-2<br>(AB_2091523) | 1:50 |
| α-Rab11 (rabbit) | Andrew lab (John Hopkins University) | | 1:500 |
| α-Nuf (guinea pig) | Sotillos lab (CABD) | | 1:500 |
| α-acetylated α-tubulin (mouse) | Invitrogen | 32-2700<br>(AB_2533073) | 1:1000 |
| α-tyrosinated α-tubulin (rat) | Invitrogen | MA1-80017<br>(AB_2210201) | 1:1000 |
| α-GFP (chicken) | Invitrogen | A10262<br>(AB_2534023) | 1:500 |
| α-β-galactosidase<br>(rabbit) | Invitrogen | A11132<br>(AB_221539) | 1:500 |
| α-Crb (mouse) | DSHB (deposited by E.<br>Knust, Max Planck<br>Institute) | Cq4<br>(AB_528181) | 1:10 |
| α-Fog (rabbit) | N. Fuse (Fuse et al.,<br>2013) | | 1:1000 |
| α-Sec15 (guinea pig) | Bellen lab (Baylor College<br>of Medicine) | | 1:2000 |
| Alexa Fluor<br>488/568/647-coupled<br>secondary antibodies | Invitrogen |  | 1:500 |

**Table 3.**
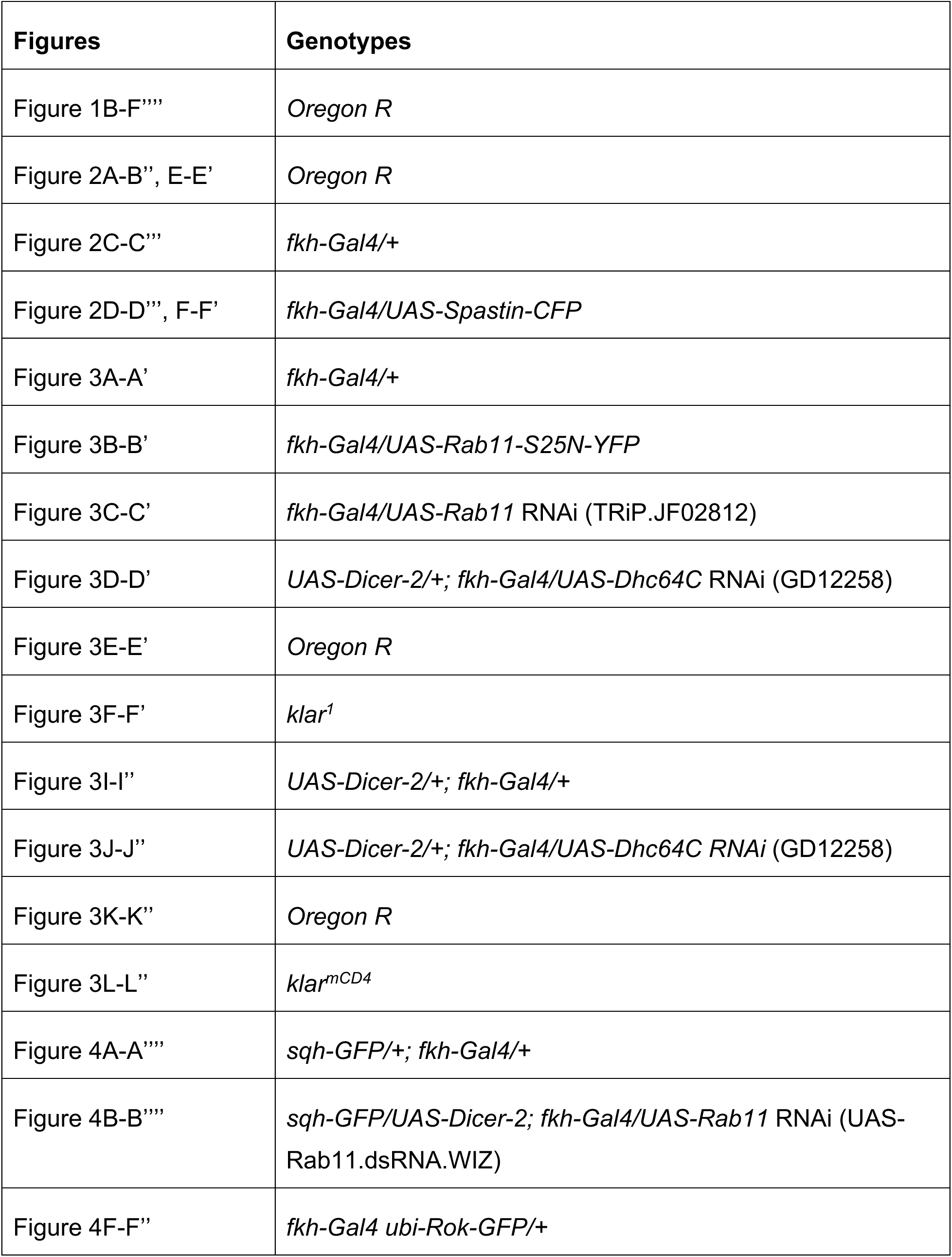
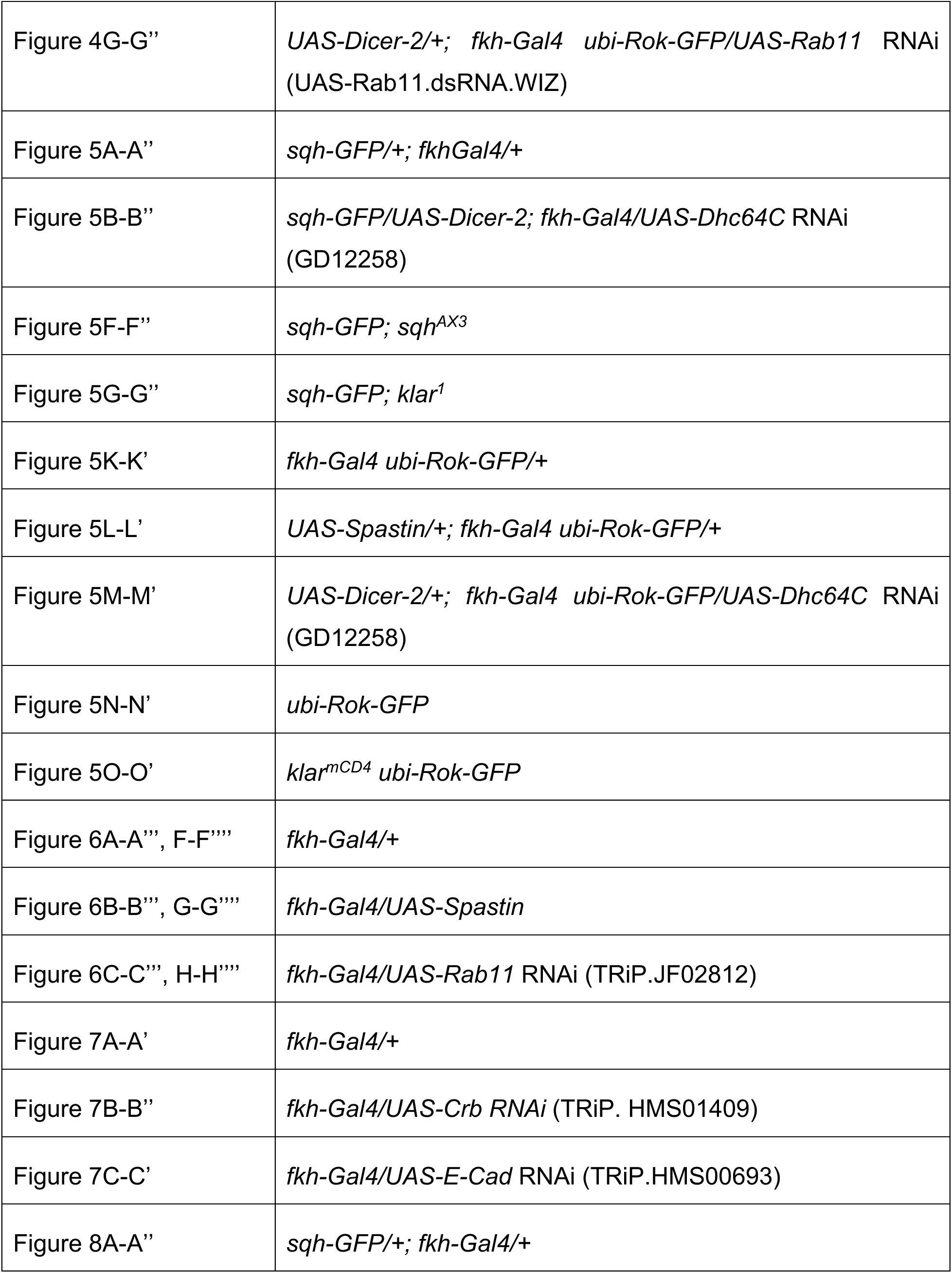
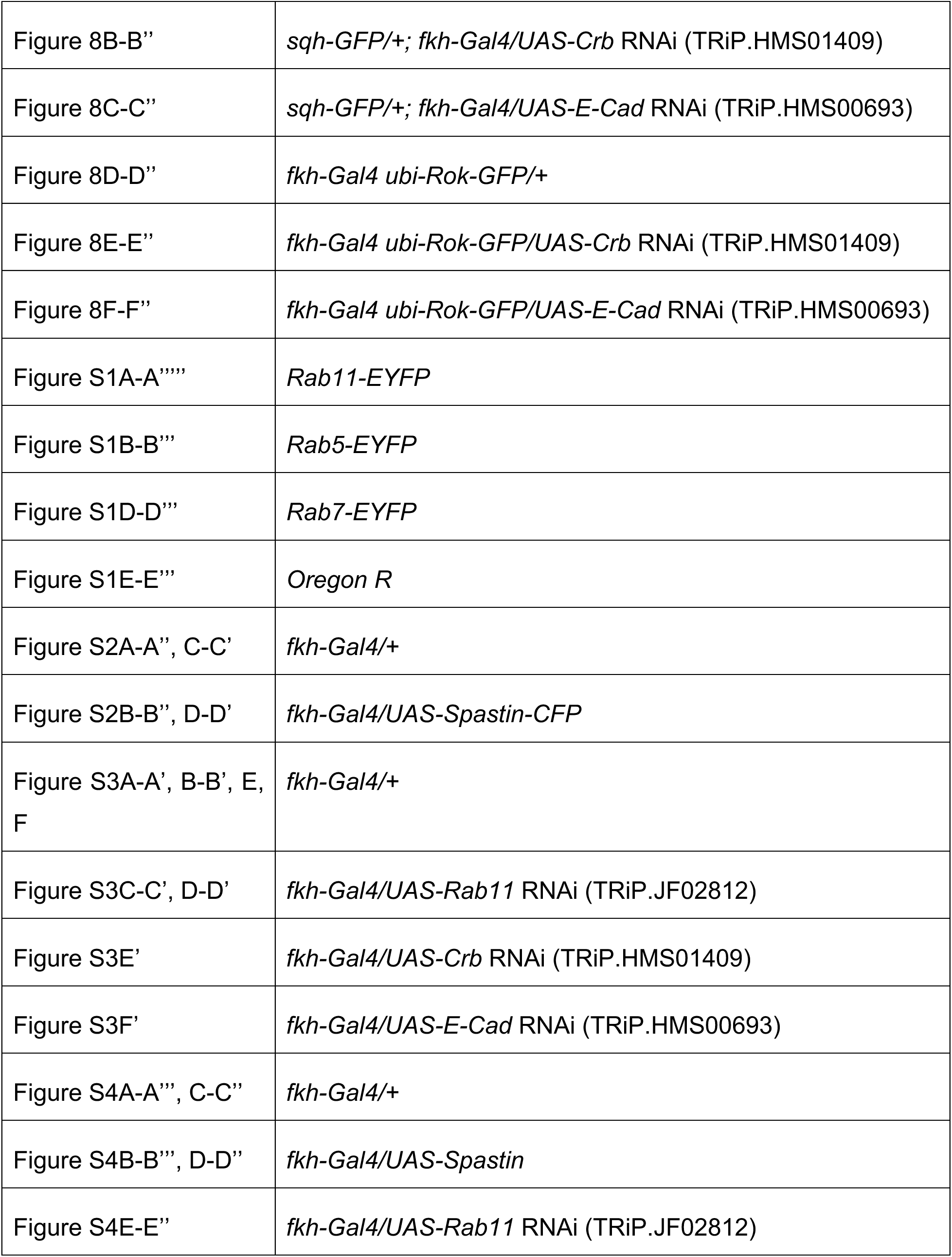
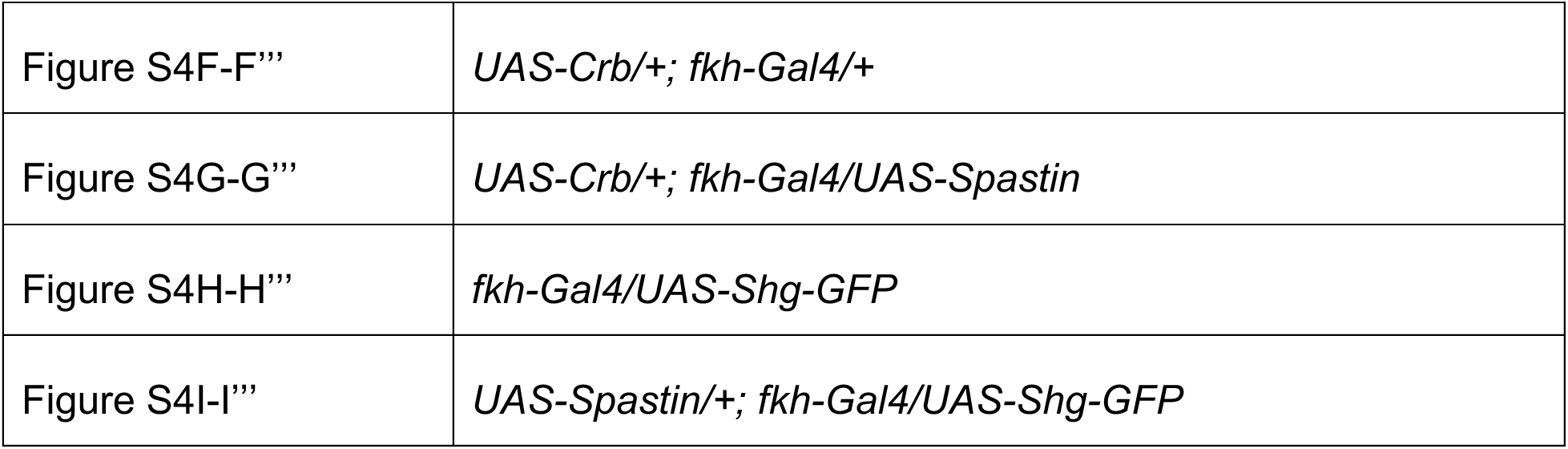
Genotypes in figure panels.

## Acknowledgments

We thank the members of the Chung laboratory for comments and suggestions. We thank A. Martin, S. Parkhurst and N. Sherwood and the Bloomington stock center for fly stocks, and D. Andrew, H. Bellen, S. Sotillos and the Developmental Studies Hybridoma Bank for antibodies. We thank Flybase for the gene information. We are grateful to A. Bohnert, C. Hanlon, A. Johnson and C. O’Kane for their helpful comments on the manuscript. This work is supported by start-up fund from Louisiana State University and the grant from the Board of Regents Research Competitiveness Subprogram GR-00005224 to S.C.

---

**Figure S1.**
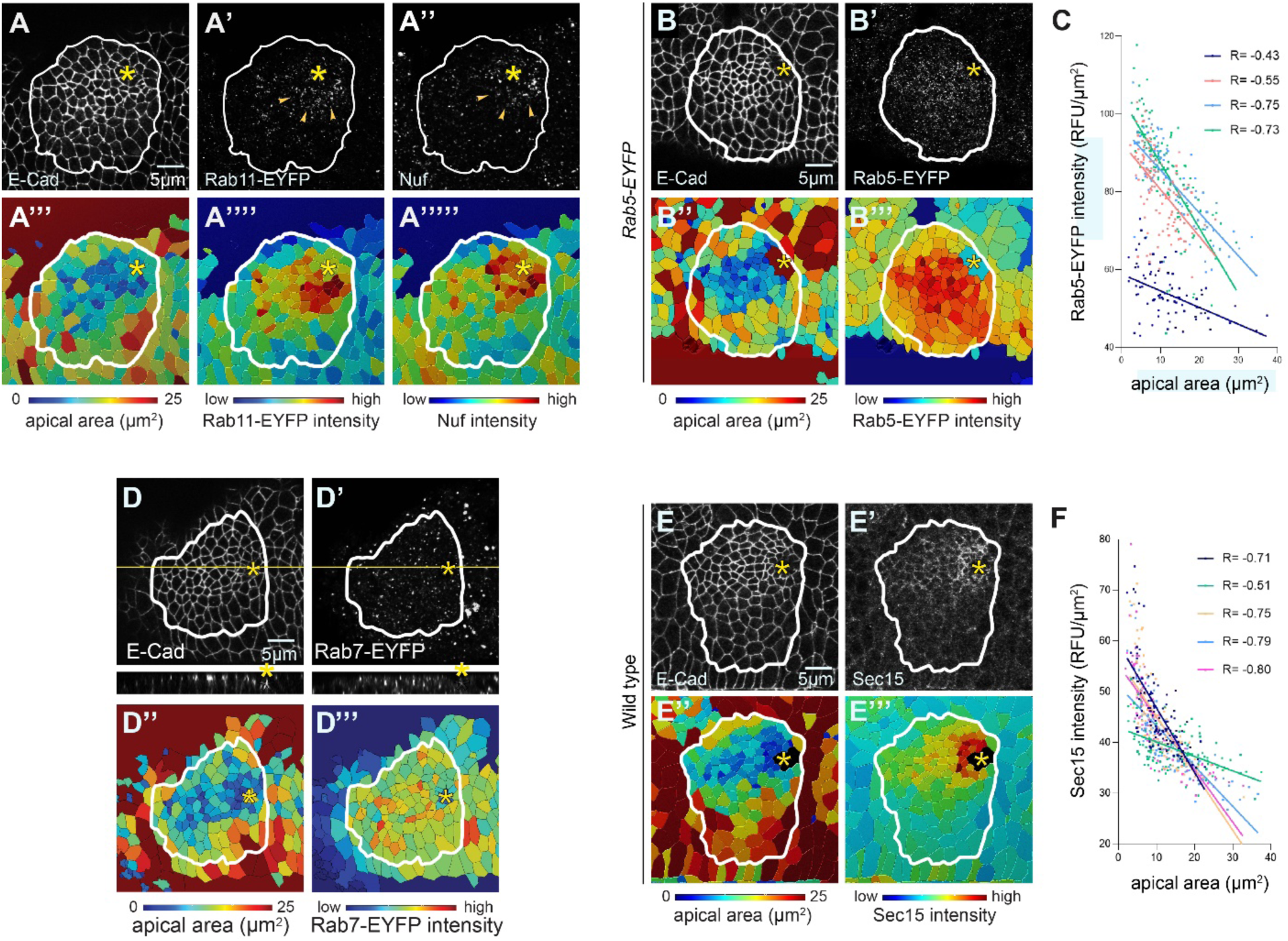
Rab11-EYFP, Rab5-EYFP and Sec15 are apically enriched in the invaginating SG. (A-A’’) *Rab11-EYFP* (an EYFP insertion at the N-terminus under the control of Rab11 regulatory sequences) SG labeled with antibodies against E-Cad (A), GFP (A’) and Nuf (A’’). (A’’’-A’’’’’) Corresponding heat maps for apical areas of SG cells (A’’’) and intensities of GFP (A’’’’) and Nuf (A’’’’’) signals. (B, B’) Confocal images of a *Rab5-EYFP* (an EYFP insertion at the N-terminus under the control of Rab5 regulatory sequences) SG stained for E-Cad (B) and GFP (B’). (B’’, B’’’) Corresponding heat maps for apical areas of SG cells (B’’) and intensity of Rab5-EYFP signals (B’’’). (C) Negative correlation between Rab5-EYFP intensity and apical areas of SG cells (n= 4 SGs; 396 cells). (D-D’) *Rab7-EYFP* (an EYFP insertion at the N-terminus under the control of Rab7 regulatory sequences) SG immunostained for E-Cad (D) and GFP (D’). (D’’, D’’’) Corresponding heat maps for apical areas of SG cells (D’’) and intensity of Rab7-EYFP (D’’’). (E-E’’’) Wild type SG immunostained for E-Cad (E) and Sec15 (E’). (E’’, E’’’) Corresponding heat maps for apical areas of SG cells (E’’) and intensity of Sec15 signals (E’’’). (F) Negative correlation between the intensity of Sec15 and apical areas of SG cells (n= 5 SGs; 527 cells). Asterisks: invagination pit. White lines, SG boundary.

**Figure S2.**
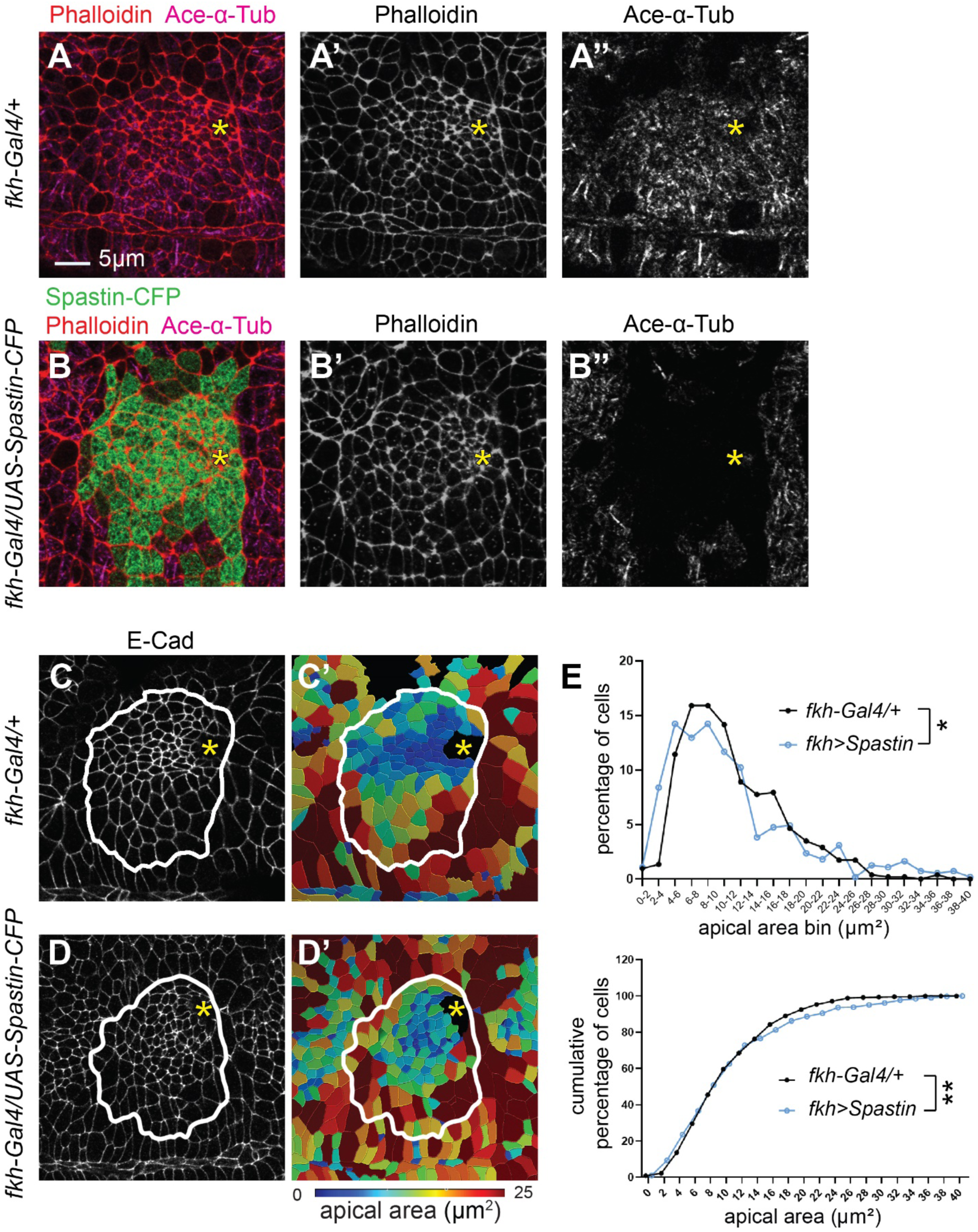
Overexpression of spastin in the SG results in loss of acetylated-*α*-tubulin. (A-B’’) Confocal images of control (A-A’’) and spastin-CFP-overexpressing (B-B’’) SGs stained for phalloidin (red) and an antibody against acetylated α-tubulin (Ace-α- Tub; magenta). Spastin-CFP (green) signals show SG-specific overexpression of spastin by *fkh-Gal4*, which leads to a loss of MTs in the SG placode. Asterisks, invagination pit. (C-D) Confocal images of controI (C) and spastin-overexpressing (D) SGs immunostained for E-Cad. (C’-D’) Corresponding heat maps for apical areas in SGs in C and D. (E) Percentage and cumulative percentage of cells with different apical areas. Mann-Whitney U test (percentage of cells) and Kolmogorov-Smirnov test (cumulative percentage of cells). n= 5 SGs (control, 517 cells; spastin overexpression, 549 cells). *, p<0.05; **, p<0.01. Asterisks: invagination pit. White lines: SG boundary.

**Figure S3.**
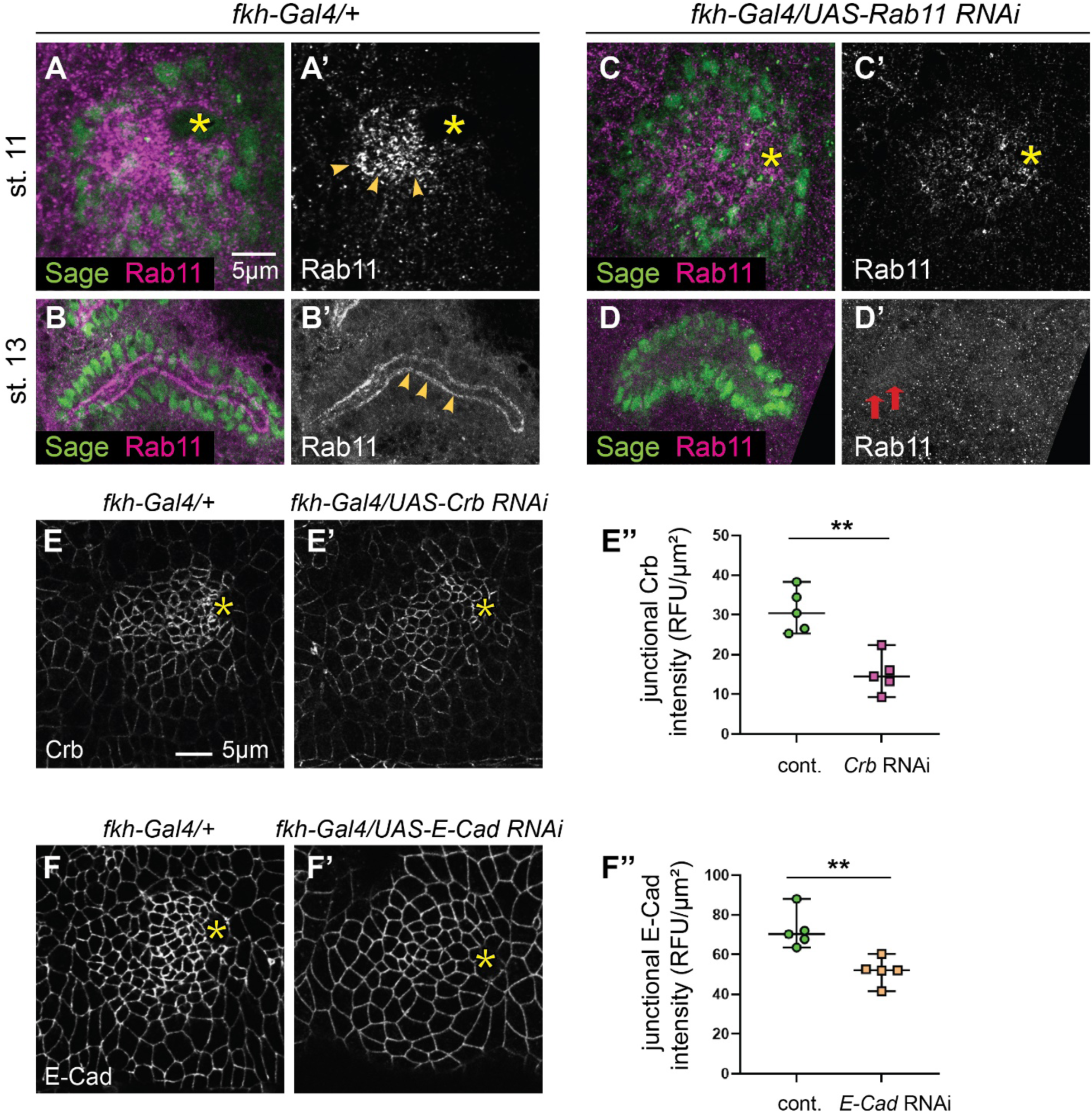
Verification of RNAi knockdown in the SG. (A-D’) Control (A-B’) and *Rab11* RNAi (C-D’) SGs immunostained for Sage (SG nuclei) and Rab11. Stage 11 (A, A’, C, C’) and stage 13 (B, B’, D, D’) SGs are shown. (E, E’) Stage 11 control (E) and *crb* RNAi (C-D’) SGs immunostained for Crb. (E’’) Quantification of the junctional intensity of Crb. (F, F’) Stage 11 control (E) and *E-Cad* RNAi (C-D’) SGs immunostained for E-Cad. (F’’) Quantification of the junctional intensity of E-Cad. n= 5 SGs for both in E’’ and F’’; 10 cells in the dorsal/posterior region of each SG. **, p< 0.01 (Welch’s t-test). Asterisks, invagination pit.

**Figure S4.**
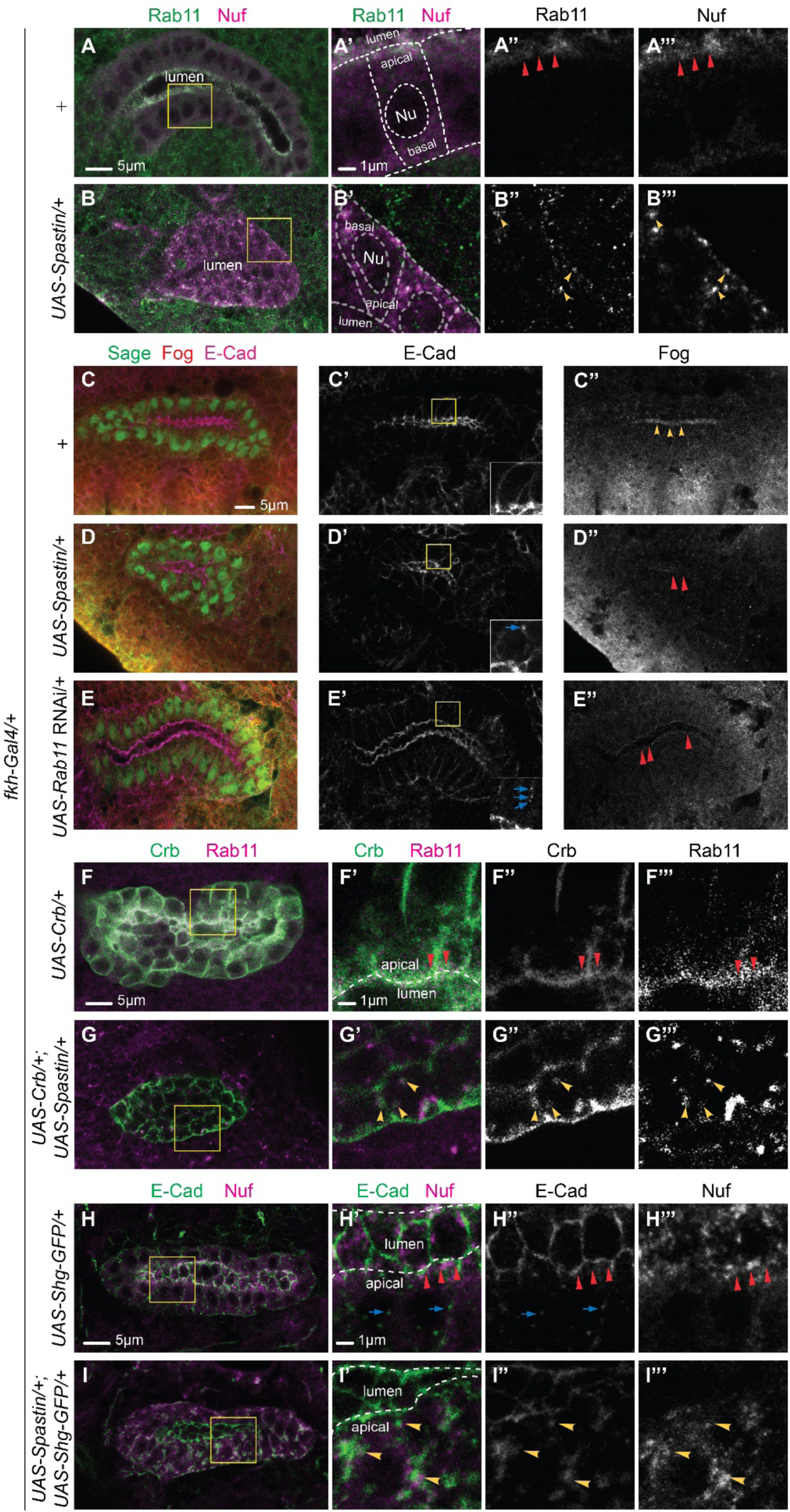
MTs play a role in apical localization of Rab11 and apical transport of Crb and E-Cad throughout SG formation. (A-B”’) Confocal images of stage 16 control (A-A’’’) and spastin-overexpressing (B-B’’’) SGs immunostained for Rab11 and Nuf. In control, Rab11 and Nuf localize in the apical region of SG cells (red arrowheads in A’’ and A’’’). In spastin-overexpressing SGs, Rab11 and Nuf are mislocalized to aggregates in the cytoplasm of cells (yellow arrowheads in B’-B’’’). (C-E’’) Confocal images of stage 13 control (C-C’’), spastin-overexpressing (D-D’’) and *Rab11* RNAi (E-E’’) SGs immunostained for Sage (SG nuclei), E-Cad and Fog. (C’-E’) Compared to strong E-Cad signals primarily at adherens junctions in the control SG (C’), strong lateral E-Cad signals are shown in spastin-overexpressing and *Rab11* RNAi SG (blue arrows in E’). Insets, higher magnification of the yellow boxed regions in C’-E’. (C’’-E’’) Compared to strong apical Fog signals in the control SG (C’’), Fog signals are faint in spastin-overexpressing (D’’) and *Rab11* RNAi (E’’) SGs. (F-G’’’) Confocal images of stage 16 SGs stained for Crb and Rab11. (F-F’’’) Overexpression of Crb causes mislocalization of Crb (green) to all membrane domains and expands membranes. Rab11 is still enriched in the apical domain (red arrowheads). (G-G’’’) Co-overexpression of spastin and Crb results in mislocalization of Crb to large aggregates in the cytoplasm, which overlap with Rab11 (yellow arrowheads). (H-I’’’) Confocal images of stage 16 SGs immunostained for E-Cad and Rab11. (H-H’’’) In the control SG, the majority of E-Cad signals are detected at adherens junctions (red arrowheads). Small punctate E-Cad signals are occasionally shown in the basolateral region (blue arrows). (I-I’’’) Co-overexpression of E-Cad and spastin causes mislocalization of E-Cad to cytoplasmic aggregates, which overlap with Nuf (yellow arrowheads). White dashed line, cell and nuclear boundaries (A’, B’) and apical membrane (F’, H’, I’). Nu, nucleus.

